# Genome-wide maps of enhancer regulation connect risk variants to disease genes

**DOI:** 10.1101/2020.09.01.278093

**Authors:** Joseph Nasser, Drew T. Bergman, Charles P. Fulco, Philine Guckelberger, Benjamin R. Doughty, Tejal A. Patwardhan, Thouis R. Jones, Tung H. Nguyen, Jacob C. Ulirsch, Heini M. Natri, Elle M. Weeks, Glen Munson, Michael Kane, Helen Y. Kang, Ang Cui, John P. Ray, Tom M. Eisenhaure, Kristy Mualim, Ryan L. Collins, Kushal Dey, Alkes L. Price, Charles B. Epstein, Anshul Kundaje, Ramnik J. Xavier, Mark J. Daly, Hailiang Huang, Hilary K. Finucane, Nir Hacohen, Eric S. Lander, Jesse M. Engreitz

## Abstract

Genome-wide association studies have now identified tens of thousands of noncoding loci associated with human diseases and complex traits, each of which could reveal insights into biological mechanisms of disease. Many of the underlying causal variants are thought to affect enhancers, but we have lacked genome-wide maps of enhancer-gene regulation to interpret such variants. We previously developed the Activity-by-Contact (ABC) Model to predict enhancer-gene connections and demonstrated that it can accurately predict the results of CRISPR perturbations across several cell types. Here, we apply this ABC Model to create enhancer-gene maps in 131 cell types and tissues, and use these maps to interpret the functions of fine-mapped GWAS variants. For inflammatory bowel disease (IBD), causal variants are >20-fold enriched in enhancers in particular cell types, and ABC outperforms other regulatory methods at connecting noncoding variants to target genes. Across 72 diseases and complex traits, ABC links 5,036 GWAS signals to 2,249 unique genes, including a class of 577 genes that appear to influence multiple phenotypes via variants in enhancers that act in different cell types. Guided by these variant-to-function maps, we show that an enhancer containing an IBD risk variant regulates the expression of *PPIF* to tune mitochondrial membrane potential. Together, our study reveals insights into principles of genome regulation, illuminates mechanisms that influence IBD, and demonstrates a generalizable strategy to connect common disease risk variants to their molecular and cellular functions.

## Introduction

Genome-wide association studies (GWAS) have now identified tens of thousands of genetic variants that influence risk for human diseases and traits^1^. Each of these associations could provide insight into biological mechanisms underlying human disease by pinpointing a particular variant, gene, and cell type that influences disease. For example, studies of protein-coding variants associated with inflammatory bowel diseases (IBD) have uncovered causal roles for *IL23R* via pro-inflammatory signaling in T cells^2^, *C1ORF106* in barrier function in enterocytes^3^, and *ATG16L1* in autophagy in a variety of immune and epithelial cells^4–7^ — thereby identifying new pathways and therapeutic targets^8^. Yet, beyond these and a handful of other examples^9–12^, connecting GWAS associations to molecular functions has proven challenging.

Three obstacles have blocked progress. One challenge is that each GWAS signal might include dozens of variants in linkage disequilibrium with one another that tag a single causal variant. The second challenge is that most causal variants for common diseases do not directly alter protein-coding sequences and instead occur in noncoding gene regulatory elements such as enhancers, which can regulate genes over long genomic distances^13,1417,18^. Finally, many enhancers appear to act in very specific cell types or cell states^15,16^, making it challenging to identify the particular cellular context in which a disease variant might act. As such, studying any given GWAS association remains daunting because it can include many possible variants, dozens of possible target genes, and dozens of possible cell types and states relevant to disease^1,19^.

Recent developments set the stage for addressing these challenges. First, toward distinguishing among multiple possible variants in a locus, recent studies have applied genetic fine-mapping methods to prioritize likely causal variants for many GWAS signals^20–23^. For example, fine-mapping identified 45 variants for IBD with at least 50% probability of causing an association signal, and 364 variants with at least 10% probability^21^. Second, we recently developed a computational approach called the Activity-by-Contact (ABC) Model to identify enhancers in a particular cell type and predict their target genes based on maps of chromatin state and 3D folding^24^. This method could allow us to build genome-wide maps of enhancer-gene regulation that describe which enhancers regulate which genes in which cell types. Together, these advances suggest a new opportunity to connect noncoding GWAS variants to their target genes and cell types.

Here, we build ABC enhancer-gene maps in 131 biosamples spanning 74 distinct cell types and tissues, and use these maps to analyze fine-mapped genetic variants associated with 72 diseases and complex traits. We find that 40% of likely causal noncoding GWAS variants (fine-mapping posterior probability >95%) overlap ABC enhancers in these cell types. For IBD, ABC links noncoding variants to 43 genes in an array of immune cell types, identifying context-specific functions for new and known genes and pathways. Together, our study provides a map of enhancer-gene regulation across cell types, demonstrates a generalizable strategy to connect thousands of risk variants to target genes, and provides a foundation for studying the molecular mechanisms that influence IBD and other common diseases.

## Results

### Genome-wide maps of enhancers and target genes in 131 biosamples

We used the ABC Model^24^ to construct genome-wide maps of enhancer-gene connections across 131 biosamples. These maps combine data from the ENCODE Consortium and other sources (in primary cell types, tissues, and immortalized or transformed cell lines) with new datasets we collected (in immune cell lines amenable to CRISPR experiments) (**Table S1**, **Table S2**, **Fig. 1a**).

**Figure 1.**
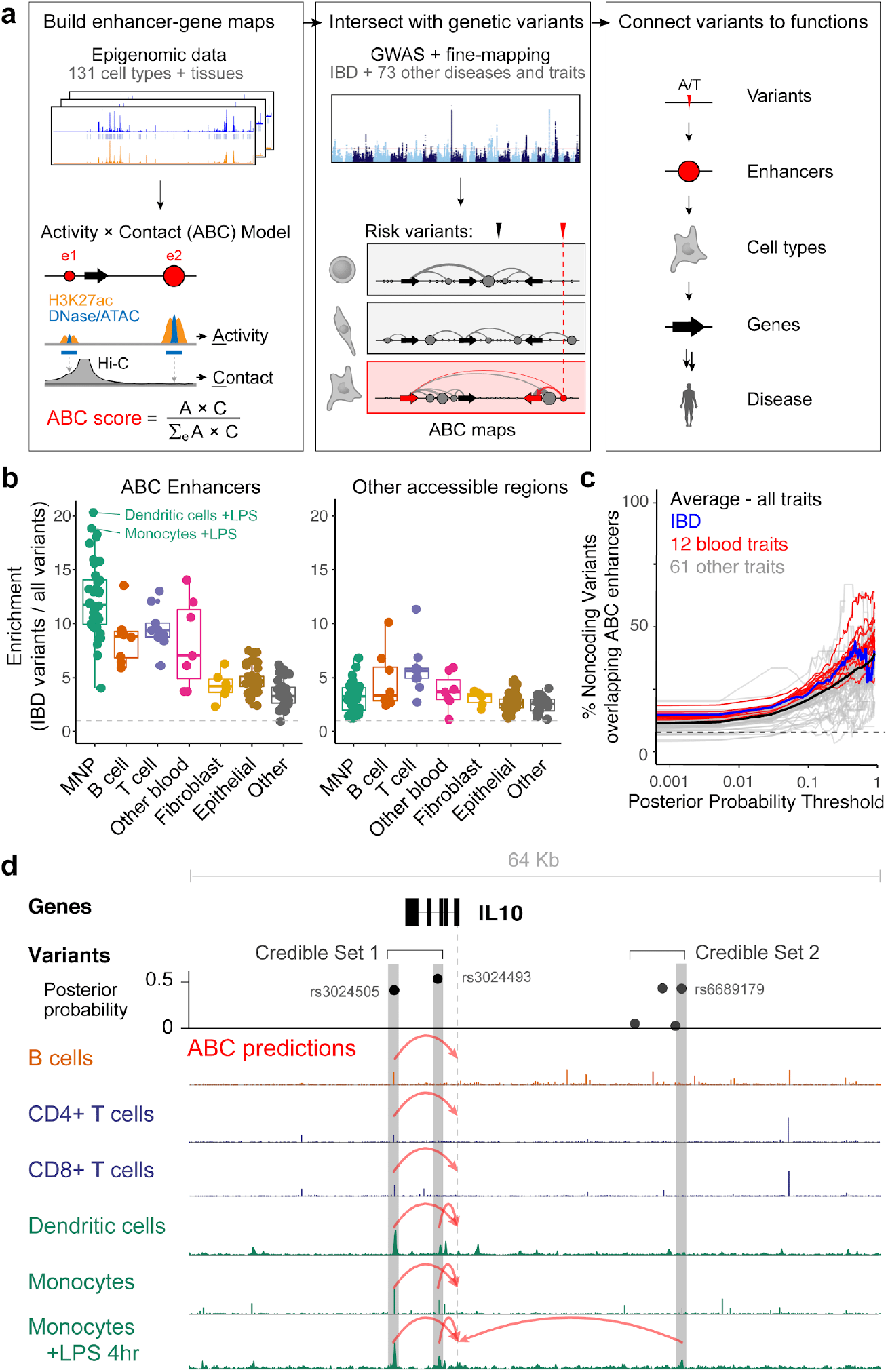
Enhancer-gene maps connect fine-mapped variants to enhancers, genes, and cell types. **(a)** Overview of approach. **(b)** Enrichment of fine-mapped IBD variants (PIP >= 10%) in ABC enhancers (left) and all other accessible regions (right) in each of 131 biosamples. ABC enhancers in mononuclear phagocytes (MNPs) including dendritic cells and monocytes stimulated with LPS show the strongest enrichments, and ABC enhancers (the subset of accessible regions predicted by ABC to regulate at least one gene) show much stronger enrichments than the other accessible regions not called as ABC enhancers. **(c)** Fraction of noncoding variants above a given PIP threshold that overlap an ABC enhancer in any biosample. Black line shows the weighted average across 72 traits. Traces are shown for PIP thresholds above which there are at least 5 variants. Dashed line shows the fraction of all common noncoding variants that overlap ABC enhancers. **(d)** ABC predictions connect two IBD GWAS signals to *IL10*. Signal tracks show DNase- or ATAC-seq (based on availability of data). Red arrows represent ABC predictions connect variants to *IL10.* Dashed line shows transcription start site (TSS). Gray bars highlight fine-mapped variants that overlap ABC enhancers in at least one cell type. Credible set 1 contains two variants, both of which overlap enhancers predicted to regulate *IL10* in various cell types. Credible set 2 contains four variants, one of which overlaps an enhancer predicted to regulate *IL10* in monocytes stimulated with LPS.

For each biosample, we defined a set of candidate elements by calling peaks in chromatin accessibility data (DNase-seq or ATAC-seq). For each candidate element and nearby active promoter, we calculated an ABC score by multiplying enhancer activity (defined in terms of chromatin accessibility and H3K27ac ChIP-seq signals in a given biosample) and element-promoter 3D contact frequency (estimated using a common Hi-C dataset averaged across 10 human biosamples) (see Methods)^24^. We defined candidate element-gene pairs that exceeded a chosen threshold on the ABC score as “enhancer-gene connections”, and elements predicted to regulate at least one gene as “ABC enhancers” (see Methods).

Across 131 biosamples, we identified 6,316,021 enhancer-gene connections for 23,219 expressed genes and 269,539 unique enhancers. In a given biosample, ABC identified an average of 48,441 enhancer-gene connections for 17,605 unique enhancers, comprising ~2.9 Mb of enhancer sequence (~12% of chromatin-accessible regions, 0.11% of the mappable genome, **Table S2**). Each enhancer regulated an average of 2.7 genes, and each gene was regulated by enhancers (**Fig. S1**). Enhancer-gene connections were very different between biosamples (on average, only 19% of enhancer-gene connections are shared between pairs of biosamples, versus 85% for biological replicates of the same biosample, see **Fig. S2**).

We compared these ABC predictions to an expanded compendium of CRISPR perturbation data including 5,755 tested distal element-gene pairs in 11 cell types and states (**Tables S3, S4**). ABC performed well at classifying regulatory connections (area under the precision-recall curve (AUPRC) = 0.64), and outperformed other methods based on assigning elements to the closest gene (AUPRC = 0.16), analyzing 3D loops or domains (best AUPRC = 0.05), correlating enhancer activity with promoter activity or gene expression (best AUPRC = 0.15), or other computational prediction methods (best AUPRC = 0.22), similar to our previous observations using a subset of this CRISPR data^24^ (**Fig. S3**, **Table S5**, see Methods).

### GWAS variants are strongly enriched in ABC enhancers

Noncoding GWAS variants are thought to regulate gene expression by affecting regulatory elements in particular cell types^13,14^, yet we have lacked accurate maps to quantify the enrichment of causal variants in enhancers. We examined how frequently genome-wide significant GWAS variants for IBD and 71 other traits overlap ABC enhancers in each biosample (**Table S6**). To overcome the challenge of identifying causal variants at each GWAS locus, we leveraged fine-mapping analyses that assigned to each variant a posterior inclusion probability (PIP) of causality and defined 95% credible sets for conditionally independent signals (here, defined as the minimal set of variants whose PIP sum to at least 95%, see Methods)^21^. To focus on higher-confidence distal regulatory signals, we examined the 24,922 fine-mapped variants with PIP ≥ 10% in credible sets that did not contain any coding or splice site variant (37-100% of credible sets for these traits, **Fig. S4a**).

The fine-mapped GWAS variants showed striking enrichments (up to 48-fold) in ABC enhancers in certain cell types, with the strongest enrichments typically occurring in cell types with biological relevance to the trait, and showed much stronger enrichments in ABC enhancers than in other chromatin accessible regions (**Fig. 1b,c**). For example, fine-mapped variants for 4 blood cell traits (monocyte, lymphocyte, erythrocyte, and platelet counts) showed strong enrichments in ABC enhancers in the corresponding cell type (11-to 16-fold compared to all variants in 1000 Genomes Version 3), and were more strongly enriched in the corresponding cell type than in other blood or unrelated cell types (**Fig. S4b**). In total, fine-mapped variants for 65 of 72 traits were significantly enriched in ABC enhancers in at least one biosample (**Table S7**).

For IBD, fine-mapped variants were significantly enriched in ABC enhancers in 65 biosamples (Bonferroni-corrected Fisher’s exact test *P* < 0.001), including 56 immune cell types/cell lines and tissue from the large intestine (**Fig. 1b**; **Table S6**). The top biosample showed 21-fold enrichment and corresponded to activated dendritic cells, which are known to play an important role in the initiation of inflammation in IBD^8,25,26^ (19 of 285 fine-mapped IBD variants (6.7%) overlapped ABC enhancers in dendritic cells, compared to 0.3% of all common variants; this included variants in 15 of 93 credible sets). Across all biosamples, fine-mapped variants were on average 2.3-fold more strongly enriched in ABC enhancers than in other chromatin accessible regions not called as enhancers by the ABC Model (two-sided signed-rank test *P* < 10^−15^, **Fig. 1b**).

We observed even stronger enrichments when we used stratified linkage disequilibrium score regression (S-LDSC) to estimate the heritability enrichment in ABC enhancers (**Table S6**). (S-LDSC considers not only variants in genome-wide significant GWAS loci but also in sub-significant loci). For example, S-LDSC found 50-fold enrichment for erythrocyte count heritability in ABC enhancers in erythroblasts and 42-fold enrichment for IBD heritability in ABC enhancers in dendritic cells.

### Enhancers likely contain a majority of causal noncoding GWAS variants

We next examined the total fraction of fine-mapped noncoding variants that overlapped ABC enhancers, as opposed to other noncoding sequences such as splice sites. Across all signals for these 72 traits, ABC enhancers contained 40% of the 2,520 noncoding variants with PIP >= 95%, compared to 7.5% of all common noncoding variants (**Fig. 1c**, **Fig. S4c**). For 12 blood cell traits, where the relevant cell types are better represented in our dataset, ABC enhancers contained 46% of 722 noncoding variants with PIP >= 95% (**Fig. 1c**). For IBD, ABC enhancers contained 4 of 10 noncoding variants with PIP ≥ 95% and 98 of 344 noncoding variants with PIP >= 10%, including variants in 54 of 93 noncoding IBD credible sets (57%).

Importantly, our analysis underestimates the proportion of fine-mapped variants residing in ABC enhancers because we still lack appropriate data for many relevant cell types. For IBD, for example, the maps are missing at least half of the >50 distinct cell types in the intestinal lamina propria^27^, including cell types known to be important for IBD (*e.g.*, Goblet and Paneth cells^8^). We anticipate that, when ABC maps are expanded to include hundreds of additional cell types, a majority of causal noncoding GWAS variants will reside in ABC enhancers (**Fig. S4d**).

### ABC maps connect GWAS variants to disease genes

Variants in enhancers influence disease through effects on the expression of nearby genes. Detailed experimental studies have demonstrated that individual noncoding variants can affect the regulation of one or more genes over long distances^10,11,28^. Yet, it has proven challenging to accurately measure or predict which gene(s) are regulated by a given variant, and which effects are relevant to disease^29–33^.

We tested how well ABC enhancer-gene maps could connect noncoding GWAS signals to target genes for IBD (**Fig. 1a**, **Note S1**). For each credible set for a trait, we intersected fine-mapped variants (PIP ≥ 10%) with ABC enhancers in biosamples globally enriched for such overlaps, and assigned the credible set to the target gene(s) predicted by ABC. In cases where multiple variants overlapped enhancers or an enhancer regulated multiple genes, we assigned the credible set to the single gene with the highest ABC score (“ABC-Max”). For example, **Fig. 1d** depicts two independent IBD credible sets in the *1q32.1* locus. Both credible sets include noncoding variants with PIP ≥ 10% that overlap ABC enhancers in monocytes stimulated with bacterial lipopolysaccharide (LPS), the biosample with the second highest enrichment for IBD (**Fig. 1c**). For both credible sets, ABC-Max predicted that these enhancers regulate multiple genes in the locus, but the gene with the highest ABC score was *IL10*, a key anti-inflammatory cytokine known to be important for IBD^8^ (**Fig. 1d**, **Fig. S5a**). Notably, ABC identified *IL10* but not two neighboring genes in the same interleukin family, *IL19* and *IL20* (**Fig. S5a**).

To systematically evaluate the predictions of ABC-Max and benchmark it against other approaches, we first examined a curated set of 64 known IBD genes previously identified based on associated coding variants or evidence regarding the effect of the gene in experimental models^8^ (see Methods, **Table S8**). We analyzed the 37 noncoding credible sets within 1 Mb of exactly one of these genes, and tested how often ABC-Max or other methods identified the known gene and distinguished it from the other genes within 1 Mb (median: 13 genes; range: 3–66). We visualized performance using a precision-recall plot, where recall is the fraction of credible sets for which the known gene is identified (sensitivity), and precision is the fraction of predicted genes corresponding to known genes (positive predictive value) (**Fig. 2a**).

**Fig. 2.**
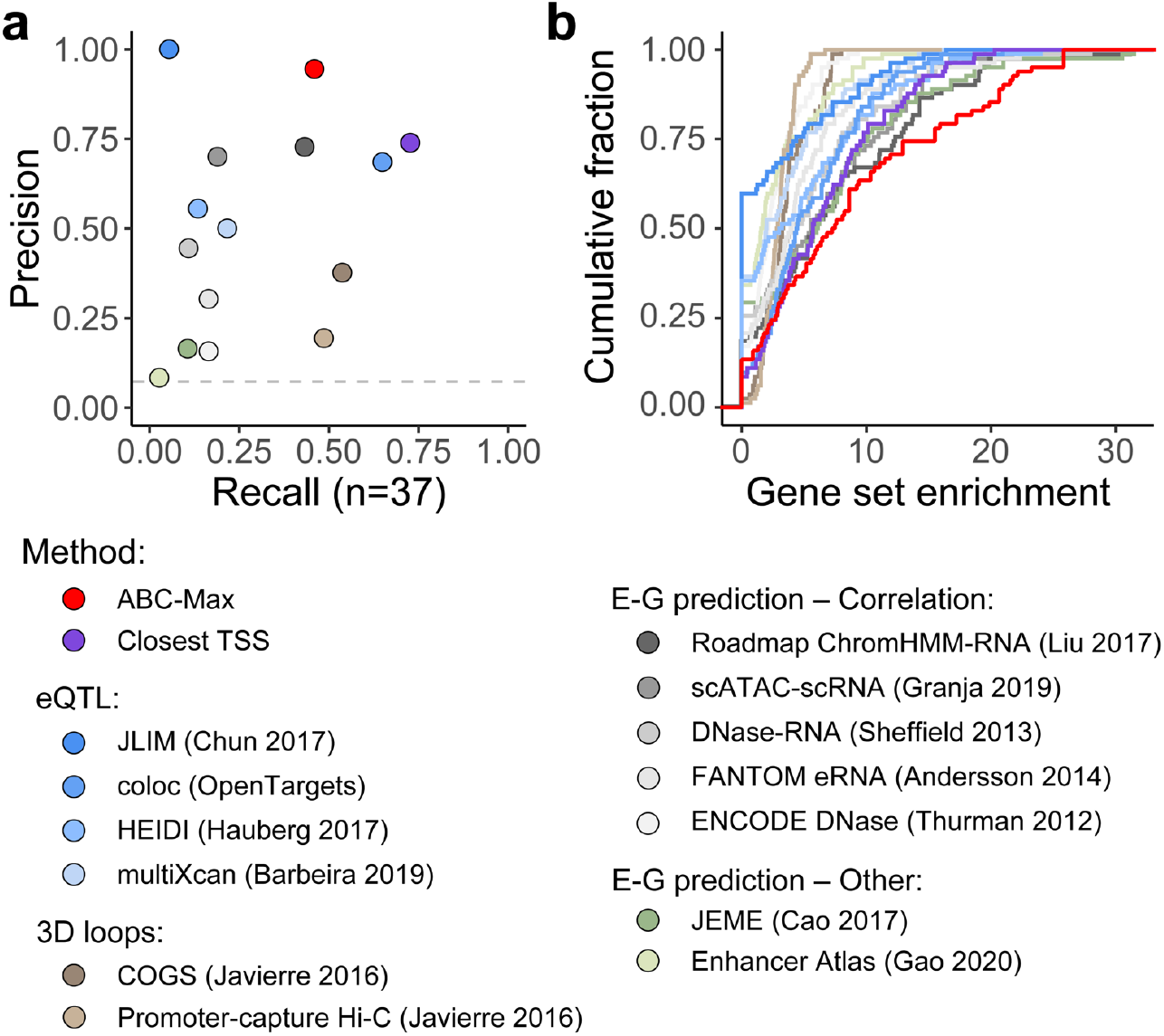
ABC enhancer maps identify known genes for IBD. **(a)** Precision versus recall for methods to connect noncoding IBD credible sets to known IBD disease genes^8^, considering 37 credible sets with exactly 1 known gene within 1 Mb. Precision = fraction of identified genes corresponding to known genes. Recall = fraction of the 37 known genes identified. See Methods for details on approaches. Where quantitative scores were available (*e.g.*, colocalization probability for eQTL coloc), plot presents the performance of choosing the gene with the best score per locus, which outperforms selecting all genes above a global threshold (see **Fig. S5b**). **(b)** Cumulative density plot showing enrichment for gene sets in MSigDB among the genes prioritized by each method^46^. For each method, we first identified the top 5 most enriched significant gene sets in the predictions of that method (82 gene sets total). Then, we calculated the levels of enrichment of all 82 gene sets in the predictions of each method.

As a baseline, we tested the simple heuristic of assigning each GWAS credible set to the closest gene. This nearest-gene method does not define a regulatory link from variant to enhancer to gene and cell type, but has been shown to assign ~70% of metabolite GWAS loci to genes with plausible biochemical functions^34^. In our IBD dataset, connecting the lead variant for each of the 37 IBD credible sets to the closest gene transcription start site (TSS) correctly identified the known IBD gene in 27 of 37 cases (73% precision, **Fig. 2a**).

We next evaluated previous methods or predictions to connect regulatory variants to disease genes, including eQTL colocalization^35,36^, transcriptome-wide association studies32, Mendelian Randomization37, promoter-capture Hi-C27, and alternative enhancer-gene prediction methods^16,38–44^ (see Methods). For nearly all methods, choosing the gene with the best score in each locus, rather than choosing all genes above the global threshold previously reported by these studies, obtained much higher precision (**Fig. S5b**). Still, most of these methods performed poorly (**Fig. 2a**). One eQTL analysis^35^ achieved 100% precision, but made predictions in only 2 loci (2 of 2 correct). The enhancer-gene prediction method with the highest precision (linking enhancers to target genes based on correlating chromatin state with gene expression across cell types from the Roadmap Epigenomics Project^38,45^) made 22 predictions and identified the known IBD genes in 16 of 22 cases (72% precision). (However, this correlation method performed much less well at predicting the results of CRISPR perturbations (AUPRC < 0.15, **Fig. S3**)).

Finally, we evaluated the predictions of ABC-Max. Of the 37 credible sets, 18 included a variant that overlapped an ABC enhancer in an enriched biosample, and ABC-Max identified the known gene in 17 of 18 cases (94% precision) (**Fig. 2a**). Thus, ABC-Max identifies a high-confidence set of genes at these IBD GWAS loci, with higher precision than other enhancer prediction methods. The fraction of loci with a prediction (recall = 49%) would likely increase upon expanding the ABC maps to include additional relevant cell types in the gut.

We performed a second analysis to compare ABC-Max to other methods for identifying genes for IBD. We analyzed all 93 fine-mapped noncoding credible sets (regardless of whether they contained a known IBD gene) and examined the extent to which the genes nominated by each method were enriched in gene sets from the Molecular Signatures Database (see Methods)^46^. Genes identified by ABC-Max for IBD showed stronger gene-set enrichments than the genes identified by other regulatory methods or by closest gene (**Fig. 2b**).

Toward the goal of understanding the good performance of ABC-Max, we made two observations. First, assigning each credible set to the gene with the strongest ABC score (“ABC-Max”; precision = 94% for known IBD genes) performed far better than assigning each credible set to all genes linked to an IBD variant (“ABC-All”; precision = 17%) (**Fig. S5b**). This was because individual noncoding variants often overlapped ABC enhancers that were predicted to regulate multiple genes (median: 3, range: 1-17), with the known gene having the highest ABC score (*e.g.*, **Fig. S5a**). This complexity appears to be a fundamental feature of mammalian gene regulation: *cis*-eQTL studies indicate that noncoding variants often regulate multiple genes in a given cell type^47^, and CRISPR experiments have identified individual enhancers that regulate up to 8 genes in *cis*^24,48^. Our observations are consistent with the idea that, while variants often affect the expression of multiple genes, only a subset of these effects are likely relevant to disease^31^.

Second, the recall of ABC-Max depended on the inclusion of relevant cell types in the analysis. To illustrate this, we repeated our ABC predictions after removing all immune cell types and gut tissue samples from our compendium. With this more limited set of samples (65 of 131 total), the recall of ABC-Max dropped from 46% to 24% (**Fig. S5b**). (The 9 genes correctly identified in this reduced dataset all appeared to be explained by variants in enhancers that were active in multiple cell types — for example, in both gut tissue and in adrenal gland, another epithelial tissue.) This highlights one feature of ABC, relative to eQTL approaches: due to the modest data requirements of ABC (chromatin accessibility and H3K27ac ChIP-seq data from a single individual), it is possible to explore a broad array of cell types and states relevant to disease.

Together, these analyses demonstrate that ABC maps can accurately connect fine-mapped variants to target genes for a substantial fraction of IBD GWAS signals, and thereby provide hypotheses for regulatory mechanisms linking causal variants to enhancers to cell types to genes.

### Context-specific functions for IBD genes and pathways

We next explored how applying ABC maps to many GWAS loci for a disease might identify genes and pathways that act in specific cellular contexts. To do so, we examined the 47 predictions made by ABC-Max for noncoding IBD credible sets, which nominated 43 unique IBD genes (**Fig. 3a**, **Table S9**). The distance from the noncoding variant in the ABC enhancer to the TSS of the predicted target gene ranged from 73 bp to 804 Kb (median: 18 Kb), and 11 of 47 predictions involved a gene that was not the closest (**Fig. 3a,b**).

**Figure 3.**
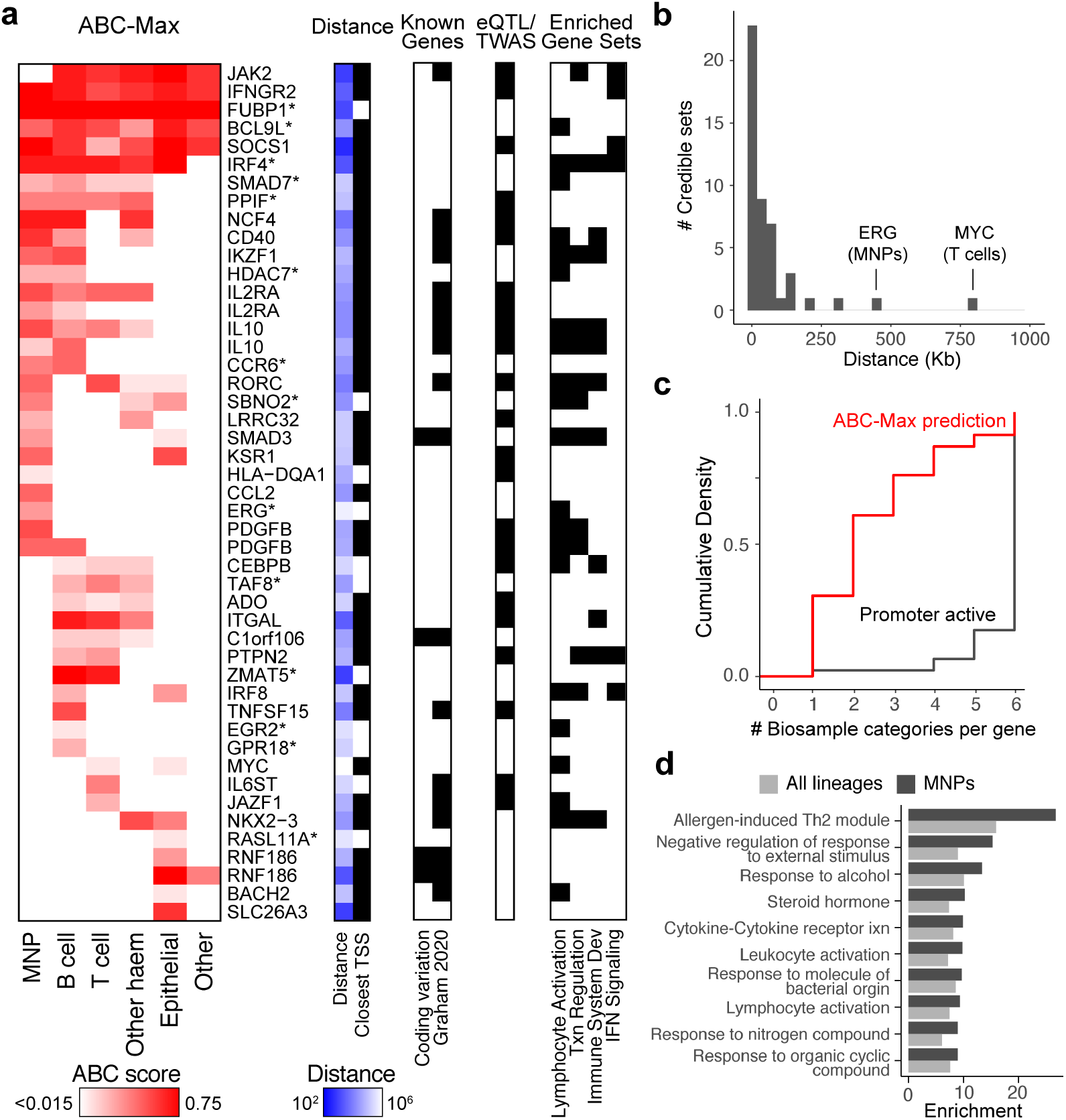
Prioritized genes for IBD. **(a)** Genes prioritized by ABC-Max for 47 noncoding IBD credible sets linking to 43 unique genes. Heatmap shows ABC scores in 6 biosample categories (maximum value across biosamples within each category). Color scale: quantiles on the ABC score, over the range [0.015, 0.75]. Blue boxes indicate log10 genomic distance from variant to gene TSS. Black boxes at right indicate if the gene has been implicated in IBD risk based on coding variants or experimental evidence about gene function^8^, is identified by prior eQTL colocalization or TWAS analyses, or is a member of selected gene sets (see Methods). Asterisks mark newly prioritized genes. **(b)** Histogram of the distances from the predicted IBD variant to the TSS of the target gene. The two genes with the greatest distances are labeled, along with the cell type of their ABC predictions. **(c)** A comparison of the number of biosample categories (cell type lineages) in which the gene promoter is active versus the number of categories in which a variant is predicted to regulate the gene by ABC-Max. **(d)** MSigDB gene sets identified when performing enrichment analysis among the genes prioritized in mononuclear phagocytes (MNPs). The enrichment for a given gene is calculated as the ratio of the frequency at which ABC-predicted genes belong to the gene set, compared to the frequency at which all genes within 1 Mb of these loci belong to the gene set (see Methods). Dark gray bars show enrichment for genes prioritized in MNPs; light gray bars show enrichment among genes prioritized in any lineage.

ABC-Max identified both known and novel IBD genes. Among the 43 prioritized IBD genes, many have previously reported functions in immunity and inflammation, and were enriched for genes in the interferon gamma pathway (6 genes; 12-fold enrichment), lymphocyte activation (11 genes; 7-fold enrichment), and regulation of transcription from RNA polymerase II promoter (21 genes; 5-fold enrichment), among other gene sets (**Fig. 3a**). For at least 26 of these genes, knockouts or other genetic models have been shown to affect the severity of experimental colitis in mice (**Table S10**). Yet, only 15 of the prioritized IBD genes were previously reported as implicated in IBD risk in humans^8,49^, and only 20 have been previously linked to IBD risk variants in large eQTL or TWAS studies (**Fig. 3a**, **Table S8**)^32,35–37,50^. The 14 newly prioritized genes included *PPIF*, a gene involved in mitochondrial metabolism whose function in immune cells is not well understood (see below).

The cell-type specificity of ABC enhancer-gene connections allowed us to categorize GWAS loci by predicted cell type(s), often identifying a cellular context more specific than the expression pattern of the gene itself (**Fig. 3a,c, Fig. S6**). For example, *IL6ST* is expressed in most cell types, but the linked IBD risk variant (rs7731626, PIP = 28%) overlapped an enhancer that was active only in T cell subsets and fetal thymus tissue (**Fig. S6d**). To validate this prediction, we used CRISPRi-FlowFISH^24^ to test whether the predicted enhancer regulates *IL6ST* expression in four immune cell lines representative of T cell, B cell, and monocytic lineages. Although *IL6ST* was expressed in each cell line, the enhancer regulated *IL6ST* expression only in a T cell line (Jurkat), and not in B cell or monocytic cell lines (GM12878, BJAB, and THP1) (**Fig. S6d**). Together, these data suggest that this IBD risk variant affects gene expression in T cells, and not in other cell types. More generally, for the 47 ABC-Max predictions, the promoter of the target gene was active in a median of 6 of 6 biosample/cell type categories (*e.g.*, T cells, B cells, see **Fig. 3a**), but the enhancer was predicted to regulate that gene in a median of only 2 categories (**Fig. 3c**, two-sided rank-sum test *P* < 10^−16^).

Connecting GWAS loci to effects in particular cell types could improve the identification of gene pathways relevant to disease. For example, the 23 IBD genes predicted in mononuclear phagocytes (MNPs: monocytes, macrophages, and dendritic cells) included a cascade of known cell-surface receptors and co-stimulatory molecules (*CD40, HLA-DQA1*, *IFNGR2*, *IL2RA*, *CCR6*), signal transduction regulators (*KSR1*, *SOCS1*), transcriptional regulators (including *IKZF1*, *ERG*, *IRF4*, *RORC*), and downstream secreted immunomodulators (*PDGFB*, *CCL2*, *IL10*). Overall, the 23 IBD genes predicted by ABC in MNPs showed much stronger gene-set enrichments in the Molecular Signatures Database (see Methods) than did the 43 IBD genes predicted without regard to cell type (up to 27-vs 16-fold enrichment, respectively; two-sided rank-sum *P* < 0.002, **Fig. 3d**), supporting the idea that grouping genes by relevant cell type can facilitate the identification of gene networks relevant to disease^27,51,52^.

### ABC links 951 genes to multiple complex traits

Beyond analysis of loci for a single trait, connecting GWAS variants to target genes across multiple traits could inform the physiological functions of genes and the selection of drug targets^53–55^. Accordingly, we used ABC maps to explore the pleiotropy of variant-gene-trait connections across 72 diseases and complex traits, for which ABC-Max identified a total of 4,976 variants that overlapped enhancers linked to 2,249 unique genes (**Table S9**).

We identified 425 genes that were linked by ABC to the same variant for multiple traits, suggesting how transcriptional regulation of these genes might influence cellular and physiological functions (**Fig. 4a-b**, **Table S11**). (105 such genes fit this criterion in the subset of 37 traits with low pairwise genetic correlation, see Methods). For example, *OSGIN1* was predicted by ABC to be regulated in hepatocytes by an enhancer that contained rs4782568. This variant has fine-mapping PIP >= 10% for 14 complex traits including serum levels of proteins and metabolites produced in the liver, including alkaline phosphatase, alanine aminotransferase, albumin, C-reactive protein, ApoB, IGF1, LDL and total cholesterol, and sex hormone binding globulin (**Fig. S7a**). *OSGIN1* encodes a protein that regulates cellular growth in response to oxidative stress^56^, and its transcriptional regulation in the liver may broadly influence hepatic metabolic functions.

**Figure. 4.**
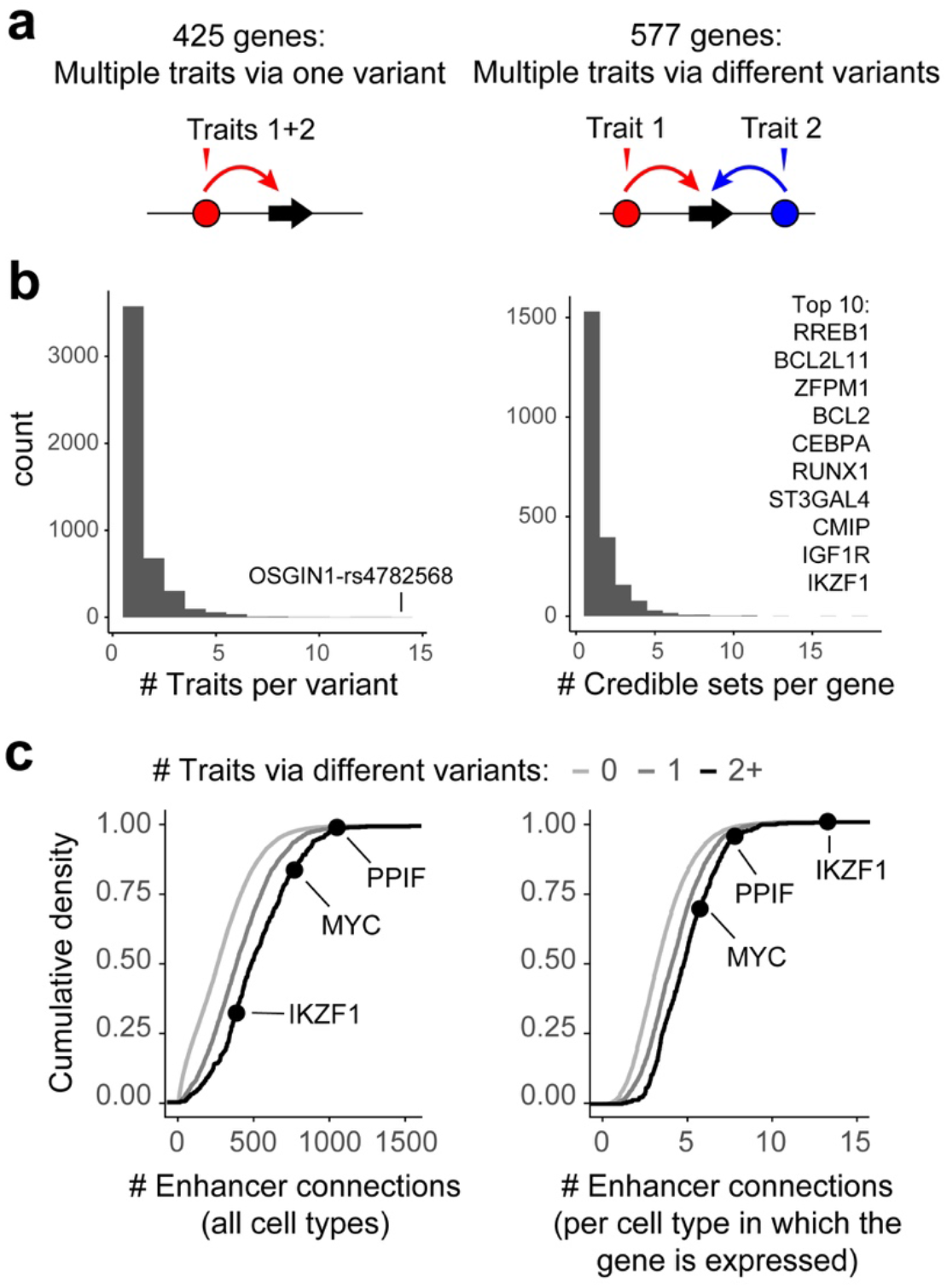
Pleiotropic effects of variants and genes across 72 diseases and complex traits. **(a)** Two classes of genes linked by ABC to multiple traits. Circles: enhancers. Black arrows: gene. Colored arrows: ABC predictions. Triangles: variants. **(b)** Histograms of properties of ABC-Max predictions, including the number of linked traits per gene, and number of GWAS signals per gene (unique credible sets with no overlapping variants with PIP >= 10%, see Methods). **(c)** Number of predicted enhancer-gene connections for genes linked by ABC to zero traits, one trait by one or more variants, or two or more traits via different variants. Three genes described in text are labeled.

We also identified 577 genes that were linked by ABC to *different* traits through *different* variants (**Fig. 4a-b**, **Table S11**; 288 genes fit this criterion in the subset of 37 traits with low genetic correlation). For each such gene, the predicted variants overlapped ABC enhancers in different sets of biosamples, indicating that cell-type specific transcriptional regulation of these genes may influence different traits. For example, *IKZF1* encodes a transcription factor involved in many stages of hematopoietic differentiation, and was linked by ABC to 12 traits via 19 unique variants in 18 credible sets, including variants associated with erythrocyte, monocyte, or neutrophil count that overlapped ABC enhancers in erythroblasts, monocytes, or CD34+ hematopoietic progenitors, respectively (**Fig. S7b**). Another such gene was *MYC*, whose expression quantitatively regulates cellular proliferation and which we previously showed has dozens of nearby GWAS associations that overlap putative enhancers^57^. Here, ABC-Max identified connections to 8 traits via fine-mapped variants, including for hematocrit in erythroblasts and lymphocyte count in B and T cells (**Table S9**).

Interestingly, these 577 genes linked to multiple traits via different variants appeared to have complex enhancer landscapes: they had (i) more predicted ABC enhancer connections (median 466 across all cell types versus 261 for other genes), (ii) more ABC enhancer connections per cell type in which the gene was expressed (median 4.8 versus 3.3) (**Fig. 4c**), and (iii) more surrounding noncoding sequence (median 301 Kb versus 128 Kb distance to the closest neighboring TSSs; this last metric is independent of ABC predictions) (**Fig. S7c**). Conversely, increased complexity of the regulatory landscape of a gene, as measured by any of these three metrics, correlated with higher odds of being linked to multiple traits via different variants (top decile: odds ratios of 2.4 to 5.5, **Fig. S7d**). These observations suggest that genes with complex enhancer landscapes are more likely to influence multiple traits, likely reflecting constraints on their precise cell-type specific transcriptional control^58^.

### Linking transcriptional regulation of *PPIF* to IBD risk

ABC predictions could accelerate experimental studies to characterize individual GWAS loci, which have been challenging due to the many possible combinations of causal variants, cell types, and target genes^1,19^. To test this, we examined the IBD risk locus at chromosome 10q22.3, where ABC predictions pointed to an unexpected gene. In this locus, the credible set contains 32 noncoding variants, all of which are located in introns of the gene *ZMIZ1*. One high-probability variant (rs1250566, PIP = 19%) overlapped an ABC enhancer in several immune cell types, including monocytes and dendritic cells (**Fig. 5a,b**). Interestingly, ABC-Max predicted that the relevant gene was not *ZMIZ1*, but rather a different nearby gene, *PPIF*. Although the variant is in an intron of *ZMIZ1*, *PPIF* has the higher ABC score because the variant is in more frequent 3D contact with the promoter of *PPIF* than with the promoter of *ZMIZ1* (by a factor of 2.3), likely because the variant is closer to the promoter of *PPIF* (61 Kb vs 218 Kb, respectively).

**Figure 5.**
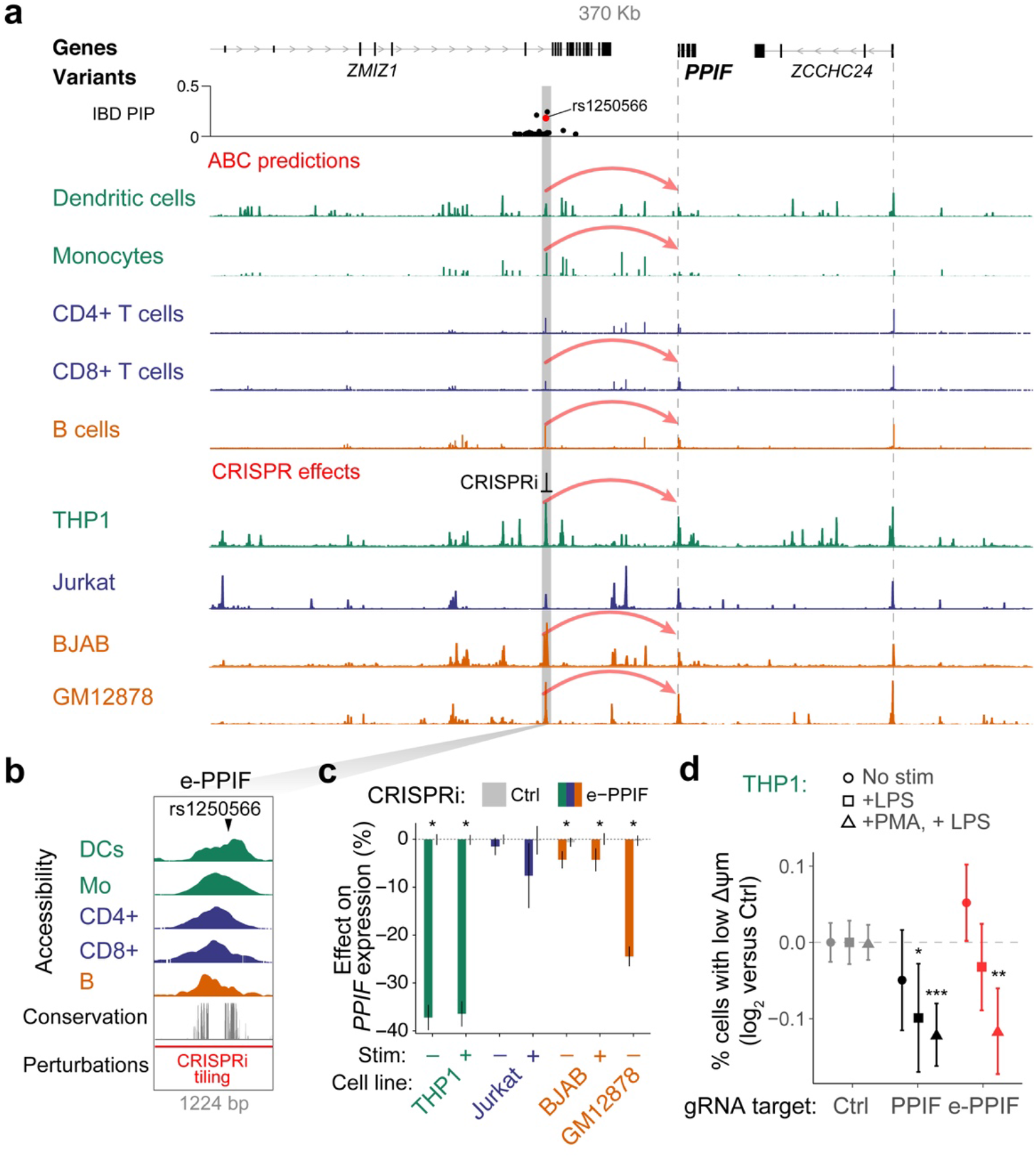
An enhancer regulates *PPIF* expression and mitochondrial function. **(a)** An IBD risk variant (rs1250566) is predicted to regulate *PPIF*. Signal tracks represent ATAC-seq or DNase-seq. Gray bar highlights enhancer containing rs1250566. Dotted lines mark TSSs. Red arcs at top indicate ABC-Max predictions connecting the enhancer to *PPIF*. Red arcs at bottom indicate that CRISPRi at the enhancer leads to a significant decrease in expression of *PPIF.* **(b)** 1224-bp region at the *PPIF* enhancer. Accessibility: DNase- or ATAC-seq from primary immune cell types (DCs=dendritic cells, Mo=monocytes). Conservation: phastCons 100-mammal alignment^68^. Red bar shows the region over which we tiled CRISPRi gRNAs. **(c)** Effects of CRISPRi at e-PPIF on the expression of *PPIF* in human immune cell lines in resting and stimulated (stim) conditions. Error bars show 95% confidence intervals of the mean. *: two-sided t-test FDR < 0.05 for 164 CRISPRi gRNAs targeting e-PPIF compared to 814 negative control (Ctrl) gRNAs. **(d)** Effects of CRISPRi gRNAs (targeting e-PPIF, PPIF promoter, or negative controls (Ctrl)) on Δψ_m_, quantified as the frequency of THP1 cells carrying those gRNAs with low versus high MitoTracker Red signal (see **Fig. S8a**). We tested THP1 cells in unstimulated conditions, stimulated with LPS, and differentiated with phorbol 12-myristate 13-acetate (PMA) and stimulated with LPS (see Methods). Error bars: 95% confidence intervals for the mean of 40, 9, and 5 gRNAs for Ctrl, PPIF, and e-PPIF, respectively. Two-sided rank-sum *P* < 0.05 (*), <0.01 (**), or <0.005 (**) versus Ctrl.

*PPIF* encodes cyclophilin D, a protein expressed in virtually all cell types that regulates metabolism, reactive oxygen species signaling, and cell death via control of the mitochondrial permeability transition pore^59–61^. *Ppif*^−/−^ knockout mice are protected from experimental colitis^62^, but *PPIF* has not been linked to risk for human IBD. *PPIF* has an extremely complex enhancer landscape: it is in the top 0.3% of genes with the most ABC enhancer-connections (1,019 across all biosamples), the top 2% with the most enhancers per biosample (average of 7.8), and the top 0.3% of genes with the most genomic sequence covered by its ABC enhancers (131 Kb) (**Fig. 4c**). The region around *PPIF* includes noncoding GWAS signals for 39 other diseases and traits, and fine-mapped variants for these phenotypes overlapped ABC enhancers in cell types including hematopoietic progenitors (for neutrophil count), B cells (for multiple sclerosis and celiac disease), and adipose tissue (for levels of insulin-like growth factor 1) (**Fig. S8a**). These observations indicate that cell-type specific transcriptional control of *PPIF* likely influences many complex traits.

To explore the enhancer landscape of *PPIF* and obtain evidence that variation in the predicted ABC enhancer could affect IBD risk, we used CRISPRi-FlowFISH^24^ to perturb each of the 163 accessible elements in a 712 Kb region around *PPIF* in four human immune cell lines, with and without stimulation with appropriate immune ligands (see Methods). We identified 14 enhancers that regulated *PPIF* expression in at least one of these cell lines and stimulated states (**Fig. S8b**, **Table S4**). The ABC enhancer that contained the fine-mapped IBD variant indeed significantly regulated *PPIF* expression in resting and activated monocytic (THP1) and B (GM12878, BJAB) cell lines, with the strongest effect in THP1 cells (37% effect on *PPIF* expression, two-sided t-test *P* < 10^−111^) (**Fig. 5c**, **Fig. S8b-d**).

Because PPIF is known to regulate the mitochondrial permeability transition pore and reduce mitochondrial membrane potential (Δψ_m_)^59^, we tested whether this *PPIF* enhancer might tune Δψ_m_ in THP1 cells. We infected cells with a pool of CRISPRi gRNAs targeting the *PPIF* enhancer and promoter, stained cells with MitoTracker Red (a fluorescent dye with higher signal in mitochondria with higher Δψ_m_), sorted cells into 3 bins based on their level of fluorescence, and sequenced the distribution of gRNAs in each bin to infer the effect of each gRNA on Δψ_m_ (**Fig. S8e**). CRISPRi targeting of the *PPIF* enhancer or promoter indeed increased Δψ_m_ in THP1 cells in LPS-stimulated, but not unstimulated, conditions (**Fig. 5d**, **Fig. S8f**), consistent with the expected direction of effect of *PPIF*. These experiments indicate that this enhancer can tune the metabolic state of mitochondria in cells responding to inflammatory stimuli.

Together, these data demonstrate that an enhancer regulates the expression of *PPIF* to influence mitochondrial function, and implicate transcriptional control of *PPIF* in genetic risk for IBD. Notably, PPIF is one of the molecular targets of cyclosporine A (CsA)^63^, an immunosuppressive drug used to treat IBD. Although the therapeutic effects of CsA have been thought to involve a different cyclophilin, *PPIA*, in T cells^64,65^, CsA also inhibits the mitochondrial permeability transition^59^ and has been observed to affect the maturation or activation of MNPs in a cell-intrinsic manner^66,67^. Our results highlight the possibility that the immunosuppressive effects of CsA could include inhibition of *PPIF*.

## Discussion

This work creates genome-wide maps including >6 million enhancer-gene connections that illuminate the functions of disease variants. These maps are based on a simple formula that involves multiplying measures of activity and contact, facilitating insights into enhancer function. These ABC enhancer-gene connections can be highly cell-type specific (**Fig. S6**), consistent with observations that noncoding variants often function in specific cell types and contexts^1,19,69^. Nearly half of enhancers are predicted to regulate multiple genes (**Fig. S1d**), matching observations from eQTL studies^19,47^. Certain genes are predicted to have up to thousands of enhancer connections across the cell types examined, and their transcriptional regulation in different cell types may influence multiple human diseases and traits (**Fig. 4c**).

At individual GWAS loci, ABC enhancer maps provide specific, actionable hypotheses linking noncoding variants to cell types, enhancers, and genes. These predictions linked 14 new genes to IBD risk and revealed a role for an enhancer of *PPIF* in tuning mitochondrial function. We have also prospectively applied ABC maps to identify and validate a variant that regulates *TET2* in hematopoietic progenitors to influence risk for clonal hematopoiesis (Bick *et al.*, in press at *Nature*^70^). By dramatically narrowing the search space of possible variants, cell types, and target genes at any given GWAS locus, ABC maps should accelerate variant-to-function studies for many diseases.

Our study has several limitations that highlight areas for future work (**Note S1**). (i) Because ABC predictions in a given cell type involve analyzing epigenomic data from a single individual, they will miss enhancers that are present only in certain genotypes or environmental states. Analyzing data across multiple individuals in healthy and disease contexts may help to capture a larger fraction of possible enhancers and interpret additional variants. (ii) While ABC maps appear to do a good job of linking noncoding GWAS variants to target genes, analyzing other complementary data — such as information about gene function — may further improve the ability to identify causal genes for common diseases. (iii) In comparing ABC-Max to other methods for identifying IBD genes (**Fig. 2**), we note that performance of each approach is a function of both the computational method and the varying cell types included in the dataset (see Methods). Generating deep eQTL maps and enhancer-gene maps in shared systems could help to directly compare various methods at identifying disease genes. (iv) Here we use bulk ATAC-seq and H3K27ac ChIP-seq to create ABC maps. Adapting this approach to leverage single-cell datasets could facilitate expanding these maps across many more cell types.

In summary, our approach illuminates a path toward creating a comprehensive map of enhancer regulation in the human genome. By refining computational models such as ABC and collecting the needed epigenomic data, it should be possible to create an accurate map of enhancers and their target genes in *cis* across thousands of cell types and states in the human body. These maps would provide a foundational reference for interpreting data from future genome-wide association studies to identify disease genes and cell types. Such a project is becoming feasible, and will be an essential resource for understanding gene regulation and the genetic basis of human diseases.

### Note S1. ABC enhancer-gene maps for interpreting the functions of noncoding GWAS variants

We set out to develop a method to build accurate enhancer-gene maps that could connect noncoding GWAS signals to target genes. To do so, we designed an approach that could (i) identify causal variants from among the many variants in LD with one another in a GWAS locus, (ii) assess the potential functions of variants on enhancers across many cell types potentially relevant to a disease, (iii) link variants in enhancers to target genes in relevant cell types, and (iv) be generally applicable to many common diseases. In the following sections, we discuss how ABC-Max combines genome-wide ABC maps of enhancer regulation with genetic maps of fine-mapped GWAS variants to accomplish these goals. We also discuss conceptual differences between ABC-Max and existing methods for linking noncoding GWAS signals to target genes.

#### ABC-Max operates on likely causal variants identified by genetic fine-mapping

A major challenge in leveraging epigenomic predictions to interpret the functions of GWAS variants has been that each GWAS locus typically contains many associated variants in linkage disequilibrium with one another, only one or several of which might be responsible for the association signal. Recent studies have begun to apply genetic fine-mapping methods to prioritize likely causal variants for many diseases and traits^20–22^. Here, we leverage results from two such studies that have applied fine-mapping to IBD and 73 complex traits, using methods that account for the possibility that there are multiple causal signals in a given locus^21^. We analyze variants with individual posterior probabilities above a threshold (>10% for most analyses), and find that these variants are up to 48-fold enriched in ABC enhancers in particular cell types. Variants above the PIP threshold that overlap ABC enhancers in enriched cell types are highly likely to be causal.

#### ABC-Max can assess enhancer-gene connections across many cell types

GWAS variants in enhancers appear to regulate gene expression in specific cell types or contexts^1,19,69^. Therefore, approaches to build regulatory maps to interpret GWAS variants must be able to assess many dozens of cell types potentially relevant to disease risk. To this end, an important feature of the ABC model is that it can make cell-type specific predictions from easily obtainable measurements of chromatin state in a given cell type (minimally, H3K27ac ChIP-seq and either ATAC-seq or DNase-seq; we previously found that cell-type specific measurements of 3D contacts are not required for optimal prediction accuracy, and so we estimate 3D contact frequencies from the average of Hi-C maps in 10 cell types^24^). As opposed to eQTL studies, which require measuring RNA expression in hundreds to thousands of individuals, ABC can make predictions based on data from a single individual. These reduced data requirements allow us to build ABC maps across 131 biosamples, leveraging a combination of existing datasets from ENCODE and Roadmap and other new epigenomic datasets that we generated here. With these maps, we can look up the potential effects of a variant across many cell types and states potentially relevant to disease.

#### ABC-Max links GWAS variants to known disease genes in relevant cell types

Noncoding GWAS variants might regulate any of the many nearby genes (5-60 within 1 Mb) and act in any of the dozens of cell types relevant to disease. To use ABC maps to prioritize among these possibilities, we first identify cell types where fine-mapped variants are globally enriched in enhancers. Second, we link fine-mapped variants that overlap enhancers in enriched cell types to the gene with the highest ABC score. This approach appears to outperform other regulatory methods at identifying genes curated in IBD GWAS loci (**Fig. 2**).

#### ABC-Max is generally applicable to many common diseases

Practically, the ABC model can be applied to many cell types and states in the human body, limited only by the ability to acquire the needed epigenomic datasets. Genetic fine-mapping will be increasingly applied to many common diseases as genetic studies in disease cohorts and biobanks expand. As such, we expect that ABC-Max will enable linking thousands of GWAS signals to relevant target genes and cell types to facilitate learning biological mechanisms of disease risk.

#### Limitations and future directions

##### ABC-Max has several limitations that highlight areas for future improvement

First, ABC-Max does not directly incorporate information about the function or sequence of a single-nucleotide variant, other than its overlap with a predicted enhancer. This affects a number of capabilities. The ABC predictions do not directly inform which transcription factors might be impacted. ABC-Max may miss cases where an activating variant creates an enhancer that does not exist in the input epigenomic datasets (due to the biosamples from which epigenomic data are derived not containing the risk haplotype). Although the ABC model appears to work well for identifying the target genes of fine-mapped GWAS variants (where we have strong evidence from human genetics that a particular variant is functional in some context), further work will be required to assess the extent to which this will help to interpret the functions of rare, *de novo*, or other variants, such as from whole-genome sequencing studies. These limitations could be addressed by combining ABC-Max with sequence-based models that can infer the effects of single-nucleotide changes on chromatin state^71,72^.

Second, ABC-Max involves prioritizing the single gene in a locus with the highest ABC score, and does not consider whether other genes also regulated by a variant might contribute to disease risk. Indeed, in a few cases, it appears that genes that do not have the highest ABC score in a locus appear to be relevant to disease (*e.g.*, *SORT1*^9^, **Table S9**), and in several loci there are multiple genes involved in relevant pathways linked to a single variant and enhancer (*e.g.*, *TNFSF15* and *TNFSF8*, **Table S8**). Future work will explore combining the regulatory predictions of ABC with computational or experimental approaches that consider gene pathways to improve the ability of the model to identify disease genes.

Finally, ABC-Max is designed to predict the functions of variants that affect enhancers, which appear to comprise a majority of noncoding causal GWAS variants. However, some causal noncoding variants likely act through other mechanisms, such as DNA topological elements, RNA splicing, or RNA translation efficiency. Further work will be required to identify and predict the functions of such variants, and to build integrative models that consider all of these mechanisms jointly.

## Methods

### Immune cell lines

We generated epigenomic data to build the ABC Model and/or performed CRISPRi experiments in the following human immune cell lines: THP1 (monocytic-like cell line, acute monocytic leukemia), BJAB (B cell-like cell line, EBV-negative inguinal Burkitt’s lymphoma), GM12878 (EBV-immortalized lymphoblastoid cell line), U937 (monocytic-like cell line, histiocytic lymphoma), and Jurkat (T cell-like, T cell leukemia).

#### Cell culture

We maintained cells at a density between 100K and 1M per ml (250K–1M per ml for GM12878) in RPMI-1640 (Thermo Fisher Scientific, Waltham, MA) with 10% heat-inactivated FBS (15% for GM12878, HIFBS, Thermo Fisher Scientific), 2mM L-glutamine, and 100 units/ml streptomycin and 100 mg/ml penicillin by diluting cells 1:8 in fresh media every three days.

#### Stimulation conditions for ABC maps and CRISPRi experiments

We stimulated BJAB cells with 4 μg/ml anti-CD40 (Invitrogen-140409-82) and 10 μg/ml anti-IgM (Sigma-I0759) for 4 hours. We stimulated Jurkat cells with 5 μg/ml anti-CD3 (Biolegend-317315) and 100 ng/ml phorbol 12-myristate 13-acetate (PMA, Sigma-P1585) for 4 hours. We stimulated THP1 cells with 1 μg/ml bacterial lipopolysaccharide (LPS) from *E. coli* K12 (LPS-EK Invivogen tlrl-peklps) for 4 hours. We stimulated U937 cells with 200 ng/ml LPS for 4 hours.

#### Stimulation conditions for ABC maps across extended timecourse in THP1 cells

For THP1 cells, we generated epigenomic data examining a longer time-course, by stimulating with PMA (100 ng/mL) for 12 hours, then removing PMA and adding LPS (1 μg/mL) and profiling at 0, 1, 2, 6, 12, 24, 48, 72, 96, and 120 hours after addition of LPS. Because THP1 cells adhere when stimulated with PMA (changing into a more macrophage-like state), we harvested cells by taking out the media, washing twice, adding TrypLE for 5 minutes at 37°C, then supplementing with 100 μL of media, removing cells from the round-bottom plate and pelleting.

### Epigenomic profiling of immune cell lines

To build ABC maps in human immune cell lines, we generated ATAC-seq and H3K27ac ChIP-seq data in BJAB, Jurkat, THP1, and U937 cells, with and without stimulation with the ligands described above.

#### ATAC-seq

We applied ATAC-seq as previously described^73^, with modifications. Briefly, we washed 50,000 cells once with 50 μl of cold 1x PBS and added 50 μl of Nuclei Isolation EZ Lysis Buffer (SIGMA NUC101-1KT) to resuspend gently, immediately centrifuging at 500xg for 10 minutes at 4°C. The lysis buffer was decanted away from the nuclei pellet. Afterwards, we resuspended the nuclei in 100 μl of Nuclei Isolation EZ Lysis Buffer again and centrifuged at 500xG for 5 minutes at 4°C and re-decant the lysis buffer, which we found to decrease mitochondrial reads although at the cost of library complexity. We then resuspended the nuclear pellet in 50 μl of transposition reaction mix (25 μl Buffer TD, 2.5 μl TDE1 (Illumina 15028212); 7.5 μl water, 15 μl PBS, to increase salinity which we found to increase signal-to-noise) and incubated the mix at 37°C for 30 minutes in a PCR block. Immediately following the transposition reaction, we split the 50 μl reaction volume into two and we added 25 μl of guanidine hydrochloride (Buffer PB, Qiagen 28606) to each as a chaotropic agent to stop the reaction and dissociate the proteins and transposase from the DNA. Keeping one of the reactions as backup, we proceeded with one by adding 1.8X SPRI beads (Agencourt A63881), waiting 5 minutes for the DNA to associate to the beads, and then washing the beads twice using 80% EtOH. We then eluted the DNA from the beads using 10 μl of water and added to it 25 μl NEBNext HiFi 2x PCR MasterMix (NEB M0541), with 2.5 uL of each of the dual-indexed Illumina Nextera primers (25 μM). We amplified the PCR reaction to 15 cycles, as previously described. We purified amplified libraries and removed adapters using two clean-ups with 1.8x volume SPRI (Agencourt A63881). We sequenced these libraries on an Illumina HiSeq 2500. We filtered, aligned, and processed the data to generate BAM files as previously described^48^.

#### H3K27ac ChIP-seq

We generated and analyzed ChIP-seq data from 5 million cells in each cell line and stimulation state, following protocols previously described^74^. Before harvesting for ChIP-seq, cells at 1 million cells per mL were replenished by a 1:2 (v/v) split in fresh media and allowed to grow for 4 hours. 10 million cells were harvested from each cell type at 500K cells/mL and washed 2x in cold PBS. Cells were resuspended in warm PBS with 1% formaldehyde (Cat #28906, Thermo Scientific) and incubated at 37°C for 10 minutes. Crosslinking was quenched by adding glycine to a concentration of 250 mM and incubating for 5 minutes at 37°C. Cells were placed on ice for 5 minutes, then washed 2x in ice-cold PBS and snap-frozen in liquid nitrogen and stored. Later, crosslinked cells were lysed in 1 mL cell lysis buffer (20 mM Tris pH 8.0, 85 mM KCl, 0.5% NP40) and incubated on ice for 10 minutes. The nuclear pellet was isolated by spinning the cell lysis mix at 5,600x*g* at 4°C for 3.5 minutes and discarding the supernatant. Nuclear pellets were lysed by adding 1 mL nuclear lysis buffer (10 mM Tris-HCl pH 7.5 ml, 1% NP-40 alternative (CAS 9016-45-9), 0.5% Na Deoxycholate, 0.1% SDS) with protease inhibitors on ice for 10 minutes. The chromatin-containing nuclear lysate was sonicated 3x using a Branson sonifier (ON 0.7s, OFF 1.3s, TIME 2 minutes, WATTS 10-12), with 1 minute rest between sonifications. Sonicated chromatin was spun down at maximum speed. 300 μL of the clarified supernatant was diluted 1:1 with ChIP dilution buffer (16.7 mM Tris-HCl pH 8.1, 1.1% Triton X-100, and 167 mM NaCl, 1.2 mM EDTA, 0.01% SDS). To immunoprecipitate H3K27ac, 3 μl of H3K27ac monoclonal antibody (Cat #39685, Active Motif) was added to each sample and rotated overnight at 4°C. The following morning, 50 uL of a 1:1 mix of Protein A (Cat #10008D, Invitrogen) and Protein G Dynabeads magnetic beads (Cat #10004D, Life Technologies) were washed with blocking buffer (PBS, 0.5% Tween20, 0.5% BSA with protease inhibitors), resuspended in 100 μl blocking buffer, and added to each sample. The samples were rotated end-over-end for 1 h at 4°C to capture antibody complexes, then washed as follows: once with 200 μl Low-Salt RIPA buffer (0.1% SDS, 1% Triton X-100, 1 mM EDTA, 20 mM Tris-HCl pH 8.1, 140 mM NaCl, 0.1% Na Deoxycholate), once with 200 μL High-Salt RIPA buffer (0.1% SDS, 1% Triton X-100, 1 mM EDTA, 20 mM Tris-HCl pH 8.1, 500 mM NaCl, 0.1% Na Deoxycholate), twice with 200 μL LiCl buffer (250 mM LiCl, 0.5% NP40, 0.5% Na Deoxycholate, 1 mM EDTA, 10 mM Tris-HCl pH 8.1), and twice with 200 μl TE buffer (10 mM Tris-HCl pH 8.0, 1 mM EDTA pH 8.0). Chromatin was then eluted from the beads with 60 μl ChIP elution buffer (10 mM Tris-HCl pH 8.0, 5 mM EDTA, 300 mM NaCl, 0.1% SDS). Crosslinking was reversed by adding 8 μL of reverse cross-linking enzyme mix (250 mM Tris-HCl pH 6.5, 62.5 mM EDTA pH 8.0, 1.25 M NaCl, 5 mg/ml Proteinase K (Cat #25530-049, Invitrogen), 62.5 μg/ml RNase A (Cat #111199150001, Roche)) to each immunoprecipitated sample, as well as to 10 μl of the sheared chromatin input for each sample brought to volume of 60 μl ChIP elution buffer. Reverse crosslinking reactions were incubated 2 h at 65°C and cleaned using Agencourt Ampure XP SPRI beads (Cat #A63880, Beckman Coulter) with a 2x bead:sample ratio. Sequencing libraries were prepared with KAPA Library Preparation kit (Cat #KK8202, KAPA Biosystems). ChIP libraries were sequenced using single-end sequencing on an Illumina Hiseq 2500 machine (Read 1: 76 cycles, Index 1: 8 cycles), to a depth of >30 million reads per ChIP sample.

### Curation of published epigenomic data

**Table S2** lists the data sources for each ABC biosample, and **Table S1** describes the epigenomic datasets generated for this study.

#### ENCODE

We downloaded BAM files for DNase-seq and H3K27ac ChIP-seq experiments from the ENCODE Portal on July 17, 2017^75^. We selected hg19-aligned BAM files that were marked as “released” by the ENCODE Portal and were not flagged as “unfiltered”, “extremely low spot score”, “extremely low read depth”, “NOT COMPLIANT”, or “insufficient read depth”.

#### Roadmap

We downloaded BAM files for DNase-seq and H3K27ac ChIP-seq from the Roadmap Epigenomics Project (http://egg2.wustl.edu/roadmap/data/byFileType/alignments/consolidated/) on July 12, 2017^45^.

#### Other studies

We downloaded FASTQ files for DNase-seq, ATAC-seq, and ChIP-seq data from 13 other studies (**Table S2**), and processed them using our custom pipelines as described below.

#### Merging cell types

We created a list of cell types across all sources for which we had at least one chromatin accessibility experiment (DNase-seq or ATAC-seq) and one H3K27ac ChIP-seq experiment. In cases where the same cell types were included in data from the Roadmap Epigenome Project and also from the ENCODE Portal, we used the processed data from Roadmap. In some cases, we combined data from multiple sources (*e.g.*, ENCODE data and our own datasets) to expand the number of cell types considered. As a result of this merging, some “cell types” in our dataset represent data from a single donor and experimental sample, whereas others involve a mixture of multiple donors and/or experimental samples.

### Processing of ATAC-seq and ChIP-seq data

We aligned reads using BWA (v0.7.17)^76^, removed PCR duplicates using the MarkDuplicates function from Picard (v1.731, http://picard.sourceforge.net), and filtered to uniquely aligning reads using samtools (MAPQ >= 30, https://github.com/samtools/samtools)^77^. The resulting BAM files were used as inputs into the ABC Model.

### Activity-by-Contact model predictions

We used the Activity-by-Contact (ABC) model (https://github.com/broadinstitute/ABC-Enhancer-Gene-Prediction) to predict enhancer-gene connections in each cell type, based on measurements of chromatin accessibility (ATAC-seq or DNase-seq) and histone modifications (H3K27ac ChIP-seq), as previously described^24^. In a given cell type, the ABC model reports an “ABC score” for each element-gene pair, where the element is within 5 Mb of the TSS of the gene.

Briefly, for each cell type, we:

1. Called peaks on the chromatin accessibility dataset using MACS2 with a lenient p-value cutoff of 0.1
2. Counted chromatin accessibility reads in each peak and retained the top 150,000 peaks with the most read counts. We then resized each of these peaks to be 500bp centered on the peak summit. To this list we added 500bp regions centered on all gene TSS’s and removed any peaks overlapping blacklisted regions (version 1 from https://sites.google.com/site/anshulkundaje/projects/blacklists)^15,78^. Any resulting overlapping peaks were merged. We call the resulting peak set *candidate elements*.
3. Calculated element Activity as the geometric mean of quantile normalized chromatin accessibility and H3K27ac ChIP-seq counts in each candidate element region. Chromatin accessibility and H3K27ac ChIP-seq signals in each candidate element were quantile normalized to the distribution observed in K562 cells.
4. Calculated element-promoter Contact using the average Hi-C signal across 10 human Hi-C datasets as described below.
5. Computed the ABC Score for each element-gene pair as the product of Activity and Contact, normalized by the product of Activity and Contact for all other elements within 5 Mb of that gene.

#### Average Hi-C

To generate a genome-wide averaged Hi-C dataset, we downloaded KR normalized Hi-C matrices for 10 human cell types^24^. For each cell type we:

1. Transformed the Hi-C matrix for each chromosome to be doubly stochastic.
2. We then replaced the entries on the diagonal of the Hi-C matrix with the maximum of its four neighboring bins.
3. We then replaced all entries of the Hi-C matrix with a value of NaN or corresponding to KR normalization factors < 0.25 with the expected contact under the power-law distribution in the cell type.
4. We then scaled the Hi-C signal for each cell type using the power-law distribution in that cell type as previously described.
5. We then computed the “average” Hi-C matrix as the arithmetic mean of the 10 cell-type specific Hi-C matrices. This Hi-C matrix (5 Kb resolution) is available here: ftp://ftp.broadinstitute.org/outgoing/lincRNA/average_hic/average_hic.v2.191020.tar.gz

#### Estimating promoter activity

In each cell type, we assign enhancers only to genes whose promoters are “active” (*i.e.*, where the gene is expressed and that promoter drives its expression). We defined active promoters as those in the top 60% of Activity (geometric mean of chromatin accessibility and H3K27ac ChIP-seq counts). We used the following set of TSSs (one per gene symbol) for ABC predictions, as previously described^24^: https://github.com/broadinstitute/ABC-Enhancer-Gene-Prediction/blob/v0.2.1/reference/RefSeqCurated.170308.bed.CollapsedGeneBounds.bed. We note that this approach does not account for cases where genes have multiple TSSs either in the same cell type or in different cell types.

For computing global statistics of ABC enhancer-gene connections (**Fig. S1**), we considered all distal element-gene connections with an ABC score >= 0.015 and within a distance of 2 Mb.

### Processing ABC predictions for variant overlaps

For intersecting ABC predictions with variants, we took the predictions from the ABC Model and applied the following additional processing steps: (i) We considered all distal element-gene connections with an ABC score >= 0.015 (see **Fig. S3**; lower threshold than our previous study^24^ to increase recall and identify gain-of-function variants that increase enhancer activity), and all distal or proximal promoter-gene connections with an ABC score >= 0.1 (based on our previous experimental data^24^). (ii) We shrunk the ~500-bp regions by 150-bp on either side, resulting in a ~200-bp region centered on the summit of the accessibility peak. This is because, while the larger region is important for counting reads in H3K27ac ChIP-seq, which occur on flanking nucleosomes, DNA sequences important for enhancer function are likely located in the central nucleosome-free region. (iii) We included enhancer-gene connections spanning up to 2 Mb.

### CRISPRi-FlowFISH

We applied CRISPRi-FlowFISH to very sensitively test the effects of distal elements on gene expression^24^. Briefly, CRISPRi-FlowFISH involves targeting putative enhancers with many independent guide RNAs (gRNAs; median = 45) in a pooled screen using CRISPR interference (CRISPRi), which alters chromatin state via recruitment of catalytically dead Cas9 fused to a KRAB effector domain. After infecting a population of cells with a gRNA lentiviral library, we estimate the expression of a gene of interest. Specifically, we: (i) use fluorescence in situ hybridization (FISH, Affymetrix PrimeFlow assay) to quantitatively label single cells according to their expression of an RNA of interest; (ii) sort labeled cells with fluorescence-activated cell sorting (FACS) into 6 bins based on RNA expression; (iii) use high-throughput sequencing to determine the frequency of gRNAs from each bin; and (iv) compare the relative abundance of gRNAs in each bin to compute the effects of gRNAs on RNA expression. CRISPRi-FlowFISH provides ~300 bp resolution to identify regulatory elements; has power to detect effects of as low as 10% on gene expression; and provides effect size estimates that match those observed in genetic deletion experiments^24^.

Here, we applied CRISPRi-FlowFISH to comprehensively test all putative enhancers in a ~700-Kb region around *PPIF*, and to validate additional selected enhancers (for 12 additional genes) that contained variants associated with IBD or other immune diseases or traits. For CRISPRi-FlowFISH experiments for *PPIF*, we designed gRNAs tiling across all accessible regions (here, defined as the union of the MACS2 narrow peaks and 250-bp regions on either side of the MACS2 summit) in the range chr10:80695001-81407220 in any of the following cell lines (+/− stimulation as described above): THP1, BJAB, Jurkat, GM12878, K562, Karpas-422, or U937. For CRISPRi-FlowFISH experiments for other genes, we included gRNAs targeting the promoter of the predicted gene and selected enhancer(s) nearby. We excluded gRNAs with low specificity scores or low-complexity sequences as previously described^24^. We generated cell lines expressing KRAB-dCas9-IRES-BFP under the control of a doxycycline-inducible promoter (Addgene #85449) and the reverse tetracycline transactivator (rtTA) and a neomycin resistance gene under the control of an EF1α promoter (ClonTech, Mountain View, CA), as previously described^57^. For each, we sorted polyclonal populations with high BFP expression upon addition of doxycycline. For GM12878 cells, we used an alternative lentiviral construct to express the rtTA with a hygromycin resistance gene, as GM12878 appeared resistant to selection with neomycin/G418.

We performed CRISPRi-FlowFISH using ThermoFisher PrimeFlow (ThermoFisher 88-18005-210) as previously described, using the probesets listed in **Table 1**. To ensure robust data, we only included probesets with twofold signal over unstained cells, and required an uncorrected knockdown at the TSS of >20%. We analyzed these data as previously described^24^. Briefly, we counted gRNAs in each bin using Bowtie^79^ to map reads to a custom index, normalized gRNA counts in each bin by library size, then used a maximum-likelihood estimation approach to compute the effect size for each gRNA. We used the limited-memory Broyden-Fletcher-Goldfarb-Shanno algorithm (implemented in the R stats4 package) to estimate the most likely log-normal distribution that would have produced the observed guide counts, and the effect size for each gRNA is the mean of its log-normal fit divided by the average of the means from all negative-control gRNAs. As previously described, we scaled the effect size of each gRNA in a screen linearly so that the strongest 20-guide window at the TSS of the target gene has an 85% effect, in order to account for non-specific probe binding in the RNA FISH assay (this is based on our observation that promoter CRISPRi typically shows 80-90% knockdown by qPCR)^24^. We averaged effect sizes of each gRNA across replicates and computed the effect size of an element as the average of all gRNAs targeting that element. We assessed significance using a two-sided t-test comparing the mean effect size of all gRNAs in a candidate element to all negative-control guides. We computed the FDR for elements using the Benjamini-Hochberg procedure and used an FDR threshold of 0.05 to call significant regulatory effects.

**Table 1.**
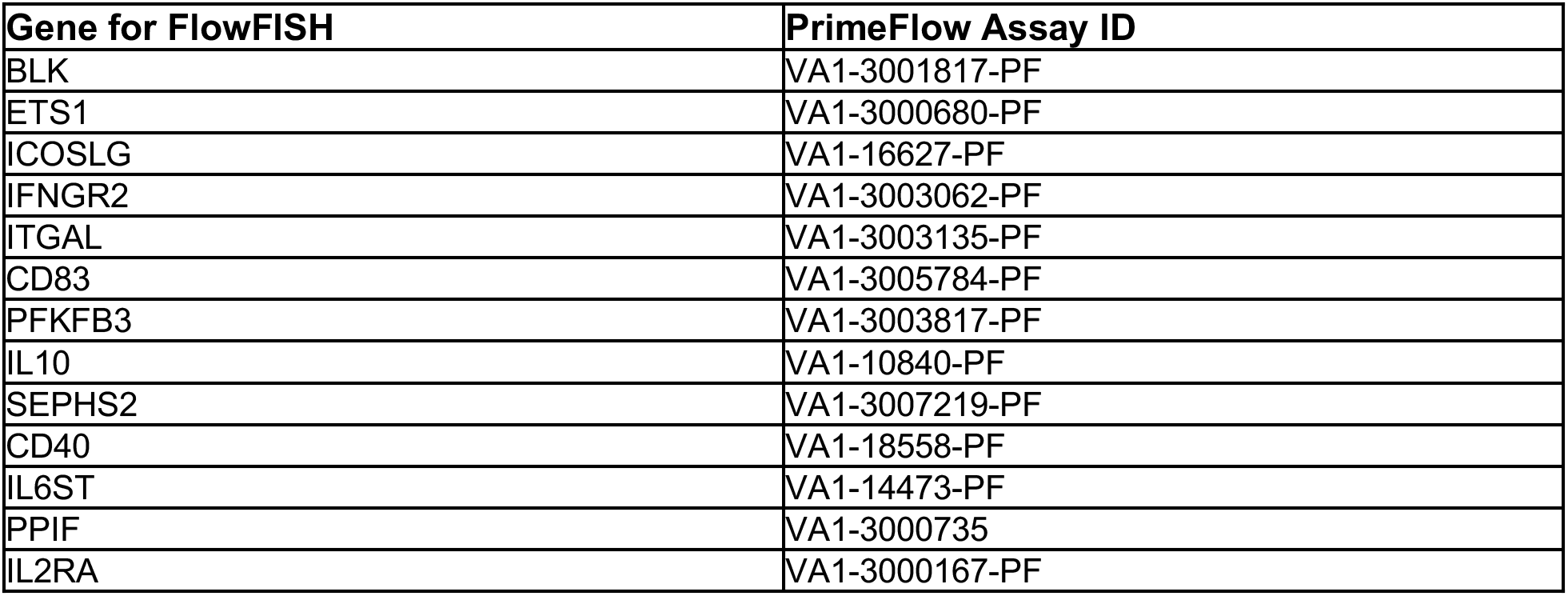
FlowFISH Probesets.

| Gene for FlowFISH | PrimeFlow Assay ID |
| --- | --- |
| BLK | VA1-3001817-PF |
| ETS1 | VA1-3000680-PF |
| ICOSLG | VA1-16627-PF |
| IFNGR2 | VA1-3003062-PF |
| ITGAL | VA1-3003135-PF |
| CD83 | VA1-3005784-PF |
| PFKFB3 | VA1-3003817-PF |
| IL10 | VA1-10840-PF |
| SEPHS2 | VA1-3007219-PF |
| CD40 | VA1-18558-PF |
| IL6ST | VA1-14473-PF |
| PPIF | VA1-3000735 |
| IL2RA | VA1-3000167-PF |

### Comparison of ABC predictions to genetic perturbations

We evaluated the ability of the ABC Score and other enhancer-gene prediction methods to predict the results of genetic perturbations using a precision recall framework. For this analysis the true positives are the experimentally measured element-gene pairs which are statistically significant and for which perturbation of the element resulted in a decrease in gene expression. For these comparisons, (i) we only considered experimentally tested elements in which the element is not within 500bp of an annotated gene transcription start site; (ii) for perturbations using CRISPRi we excluded pairs in which the element resides within the gene body of the assayed gene; (iii) we excluded non-significant pairs for which the power to detect a 25% change in gene expression was < 80%; and (iv) we only included pairs for which the gene is protein-coding (although the ABC model can make predictions for non-coding genes, many of the other predictions methods we compare to do not make predictions for such genes).

For each experimentally measured element-gene-cell-type tuple, we intersected this tuple with the tuple in the predictions database corresponding to the same cell type, same gene and overlapping element. In cases in which the genomic bounds of an experimentally tested element overlap multiple predicted elements, we aggregated the prediction scores using an aggregation metric appropriate to each individual predictor (for ABC we used ‘sum’, for correlation- or confidence-based predictors we used ‘max’). Similarly, if the predictor did not make a prediction for a particular tuple, it received an arbitrary quantitative score less than the least confident score for the predictor (for ABC we used 0, for other predictors we used 0, −1, 1 as appropriate). Table S5 lists the experimental data merged with the predictions.

In the cases in which an enhancer-gene prediction method did not make cell-type specific predictions, we evaluated the predictions against experimental data in all cell types (**Fig. S3b**). We calculated the area under the precision-recall curve (AUPRC) for predictors, or, if the predictor was defined at only one point, we multiplied the precision by the recall.

### Similarity of ABC Predictions among replicates and biosamples

We evaluated the reproducibility of ABC predictions derived from replicate epigenetic experiments. For each biosample in which independent biological replicate experiments for both ATAC-Seq (or DNase-Seq) or H3K27ac ChIP-Seq were available, we generated ABC predictions for replicates 1 and 2 separately. In order to facilitate the reproducibility analysis, when computing the ABC Scores for replicate 2, we used the candidate enhancer regions from replicate 1. (Using different sets of candidate regions can confound computing reproducibility. For example, the procedure to define candidate regions (peak calling, extending and merging) could call two separate ~500bp regions in one replicate, but merge them into a ~1-kb region in the second replicate due to minor differences in the peak summits between replicates. In such a case the ABC Score of the ~1-kb region would be equal to the sum of the ABC Scores of the 500-bp regions.)

We then evaluated the quantitative reproducibility of the predictions (**Fig. S2b**) and the number of predictions shared between replicates (**Fig. S2c**). We observed that on average 85% of enhancer-gene predictions in one replicate are shared in the other replicate (at an ABC Score threshold of 0.015). The fraction of shared connections between biological replicates increased as the ABC score cutoff increased: 95% of connections called in replicate 1 at a higher confidence threshold of 0.02 were also called in replicate 2 (at the default threshold of 0.015).

We also evaluated the extent to which the reproducibility of ABC predictions depends on the reproducibility of the underlying epigenetic data. For each biosample, we computed the correlation between the ATAC-Seq (or Dnase-Seq) or H3K27ac ChIP-Seq signals in the candidate regions for that biosample. As expected, we observed that the fraction of shared ABC predictions between replicates increased as the correlation of the underlying epigenetic data increased (**Fig. S2d**).

We used a similar calculation to compare ABC predictions across cell types and biosamples. For each pair of biosamples we computed the fraction of predicted enhancer-gene connections shared between the pair. For this analysis we used the shrunken ABC elements (~200bp, see above) and considered two connections to be shared if the elements overlapped at least 1 bp and predicted to regulate the same gene.

### Genetic data and fine-mapping

We downloaded summary statistics for IBD, Crohn’s disease (CD), and ulcerative colitis (UC) (European ancestry only)^80^ from https://www.ibdgenetics.org/downloads.html. We obtained fine-mapping posterior probabilities and credible sets from Huang et al.^21^, and analyzed the top two conditionally independent credible sets in each locus. We also analyzed variants from IBD GWAS loci that were not fine-mapped in this study^49,80^; for each such locus, we analyzed all 1000 Genomes variants in LD with the lead variant (*r*^2^ > 0.2) and weighted each variant evenly (probability = 1 / number of variants in LD). We observed similar results for cell type enrichments with or without including these non-fine-mapped sets. Throughout this text, analyses of “IBD” signals are defined as signals associated with CD, UC, or both.

We obtained fine-mapping results and summary statistics for 73 other traits based on an unpublished analysis (Jacob Ulirsch, Masahiro Kanai, and Hilary Finucane) that analyzed data from the UK Biobank (Application #31063; fine-mapping available at https://www.finucanelab.org/data). In this analysis, fine-mapping was performed using the Sum of Single Effects (SuSiE) method^81^, allowing for up to 10 causal variants in each region. Prior variance and residual variance were estimated using the default options, and single effects (potential 95% CSs) were pruned using the standard purity filter such that no pair of variants in a CS could have r^2^ > 0.25. Regions were defined for each trait as +/− 1.5 Mb around the most significantly associated variant, and overlapping regions were merged. As inputs to SuSiE, summary statistics for each region were obtained using BOLT-LMM^82^ for quantitative traits and SAIGE^83^ for binary traits, in sample dosage LD was computed using LDStore^84^, and phenotypic variance was computed empirically. Variants in the MHC region (chr6: 25-36 Mb) were excluded as were 95% CSs containing variants with fewer than 100 minor allele counts. Coding (missense and predicted loss of function) variants were annotated using the Variant Effect Predictor (VEP) version 85^85^. For analysis with ABC, we excluded neuropsychiatric traits (for which we expect existing enhancer-gene maps will not include the appropriate cell types), and analyzed only the variants that SuSIE assigned to belong to 95% credible sets (cs_id != −1).

For all traits, except where specified, we considered only the “noncoding credible sets” — *i.e.*, those that did not contain any variant in a coding sequence or within 10 bp of a splice site annotated in the RefGene database (downloaded from UCSC Genome Browser on 24/06/2017)^86^. We note that predictions for all credible sets, both coding and noncoding, are reported in **Table S9** to facilitate future analyses.

### Defining enriched biosamples for each trait

For a given trait, we intersected variants with PIP >= 10% in noncoding credible sets with ABC enhancers (or other genomic annotations). For each biosample, we calculated a *P*-value using a binomial test comparing the fraction at which PIP >= 10% variants overlapped ABC enhancers with the fraction at which all common variants overlap ABC enhancers in that cell type. We calculated the latter using common variants in 1000 Genomes as described in the S-LDSC section. For each trait, we defined a biosample as significantly enriched for that trait if the Bonferroni-corrected binomial *P*-value was < 0.001.

### Stratified linkage disequilibrium score regression (S-LDSC)

We used S-LDSC to assess the enrichment of disease or trait heritability in ABC enhancers, considering all variants across the genome^87^. We analyzed the ABC enhancer regions as defined above, and ran LD score regression using the baselineLD_v1.1 model using the 1000G_EUR_Phase3_baseline file (downloaded from https://data.broadinstitute.org/alkesgroup/LDSCORE/). For comparison, we also analyzed heritability enrichment in all other accessible regions for each trait. Specifically, we took the list of MACS2 peaks (FDR < 0.05), removed those that overlapped ABC enhancers, and used these regions in S-LDSC.

### Partitioning the genome into disjoint functional categories

To compare the frequency of variants occurring in ABC enhancers as opposed to other functional elements such as coding sequences and splice sites (**Fig. S4c**), we partitioned the genome into the following functional categories, using the RefGene database (downloaded from UCSC Genome Browser on 24/06/2017): coding sequences (CDS), 5’ and 3’ untranslated regions (UTR) of protein-coding genes, splice sites (within 10 bp of a intron-exon junction of a protein-coding gene) of protein-coding genes, promoters (±250 bp from the gene TSS) of protein-coding genes, ABC enhancers in 131 biosamples, other accessible regions in the same biosamples not called as ABC enhancers, and other intronic or intergenic regions. These categories may overlap; a disjoint annotation was created by assigning each nucleotide to the first of any overlapping categories in the order above (*e.g.*, nucleotides in both coding sequences and ABC enhancers were counted as coding sequences).

### Evaluating gene prediction methods

#### Curated genes for inflammatory bowel disease

We analyzed a list of IBD disease genes curated by Graham and Xavier (2020).^8^ To evaluate methods to connect noncoding GWAS variants to genes, we analyzed credible sets within 1 Mb of exactly 1 of these known genes that did not contain any protein-coding or splice site variants. In cases where the gene was curated based on evidence from coding variation, we examined nearby conditionally independent noncoding signals, which might act via regulatory effects on the same gene that carries the coding variant.

#### Gene set enrichment for IBD predictions

As a second approach for comparing methods for identifying causal genes in IBD GWAS loci, we examined the extent to which the predicted genes were enriched for any gene sets. To do so, we downloaded curated and Gene Ontology gene sets from the Molecular Signatures Database^46^. We analyzed all 93 noncoding IBD credible sets. For each gene set, we tested whether it was enriched in the genes predicted by a given method, using the set of all genes within 1 Mb of IBD credible sets as the background, excluding HLA genes. For **Fig. 2b**, we applied this approach to each of the methods described in **Fig. 2a**, selected the 5 gene sets with the highest enrichment that also had at least five identified genes and hypergeometric test *P*-value < 10^−4^. We plotted a CDF of the enrichments for each of the methods across the union of the top 5 gene sets identified by any of the methods.

### Comparisons to alternative variant-to-gene prediction methods

We compared ABC-Max to alternative methods to link regulatory variants to disease genes.

#### *eQTL Colocalization* (Open Targets Platform)

OpenTargets.org performed colocalization analysis between IBD GWAS signals^49,80^ and eQTLs and pQTLs using coloc. This analysis involved QTL datasets from a variety of sources including dozens of human tissues and many immune cell types. We downloaded colocalization results from ftp://ftp.ebi.ac.uk/pub/databases/opentargets/genetics/190505/v2d_coloc/ on February 1, 2020, and defined two gene sets: genes showing colocalization with an eQTL or pQTL in an immune cell type or spleen, blood, or gut tissue; and genes showing colocalization with an eQTL or pQTL in any biosample. We considered genes with coloc h4 probability >= 0.9, and h4/h3 ratio >= 2. We used the coloc h4 probability to rank genes within each locus.

#### *eQTL Colocalization* (JLIM)

Chun *et al.* tested colocalization of IBD GWAS signals with eQTLs in CD4+ T cells, CD14+ monocyte, and LCLs^35^. We obtained their colocalized genes from Table 2. We used the JLIM p-value to rank genes within each locus.

#### *TWAS* (*S-PrediXcan* and *multiXcan*)

Barbeira *et al.* developed multiXcan and compared GTEx v7 eQTLs to IBD summary statistics^32^. We downloaded Dataset 6 and compared genes within each locus using the multiXcan p-value.

#### Mendelian randomization

Hauberg *et al.* used a Mendelian randomization based approach (HEIDI) to connect IBD GWAS signals to effects on gene expression using eQTL data from 24 tissues^37^. We downloaded Table S3 and defined predicted genes using either all tissues or those observed in whole blood. We used the SMR false discovery rate to rank genes within each locus.

#### COGS

Javierre *et al.* (2016) used promoter-capture Hi-C data in many blood cell types to link GWAS variants to target genes^88^. We downloaded Table S3 (Tab 2) and analyzed genes linked with COGS scores >=0.5.

In all cases, we combined predictions of disease genes for IBD, UC, and CD.

### Comparisons to previous enhancer-gene predictions

We compared the ABC model to methods using alternative enhancer-gene linking approaches. For each of the methods below, we downloaded previous predictions of enhancer-gene links, and assessed (i) their ability to predict enhancer-gene regulation in CRISPR datasets (**Fig. S3**) and (ii) their ability to identify IBD genes (**Fig. 2**). For the latter analysis, we used the predictions from each method to overlap fine-mapped variants (PIP >= 10%) with enhancers in any cell type and assigned variants to the predicted gene(s).

#### Promoter-capture Hi-C

We downloaded Data S1 peak data from Javierre *et al.* (2016)^88^, representing promoter-capture Hi-C data from 9 hematopoietic cell types, and selected the promoter-distal region pairs with CHiCAGO score >= 5. For comparison to CRISPR data we used the CHiCAGO score as a quantitative predictor.

#### *DHS-promoter correlation* (ENCODE2)

Thurman *et al.* (2012) linked distal accessible elements with gene promoters by looking at correlation of DNase I hypersensitivity across 125 cell and tissue types from ENCODE^16^. We downloaded these links from ftp://ftp.ebi.ac.uk/pub/databases/ensembl/encode/integration_data_jan2011/byDataType/openchrom/jan2011/dhs_gene_connectivity/genomewideCorrs_above0.7_promoterPlusMinus500kb_withGeneNames_32celltypeCategories.bed8.gz. GWAS loci with high-confidence fine-mapped variants that overlapped these regions were assigned to the linked gene(s).

#### *eRNA-mRNA correlation* (FANTOM5)

Andersson *et al.* (2014) linked transcriptional activity of enhancer and transcription start sites using the FANTOM5 CAGE expression atlas^40^. We downloaded these predictions from http://enhancer.binf.ku.dk/presets/enhancer_tss_associations.bed.

#### *Enhancer-gene correlation* (Ernst Roadmap)

Liu, Ernst *et al.* (2017) correlated gene expression with five active chromatin marks (H3K27ac, H3K9ac, H3K4me1, H3K4me2, and DNase I hypersensitivity) across 56 biosamples, and then used these correlation links to make predictions for the predicted enhancers (regions with the “7Enh” ChromHMM state) in 127 biosamples from the Roadmap Epigenome Atlas^38,45^. We downloaded these predictions from www.biolchem.ucla.edu/labs/ernst/roadmaplinking and made predictions using the “confidence score”.

#### *Enhancer-gene correlation* (Granja single-cell RNA and ATAC-seq)

Granja *et al.* (2019) analyzed single-cell ATAC-seq and RNA-seq data in peripheral blood and bone marrow mononuclear cells, CD34+ bone marrow cells, and cancer cells from leukemia patients, and correlated ATAC-seq signal in distal elements with the expression of nearby genes^39^. We downloaded these predictions from https://github.com/GreenleafLab/MPAL-Single-Cell-2019, and used the correlation in healthy samples as the quantitative score. Cell-type specific links were not reported.

#### EnhancerAtlas 2.0

Gao *et al*. (2020) used *EAGLE* to predict enhancer-gene interactions across a number of human tissues and cell lines^43^. The method calculates a score based on six features obtained from the information of enhancers and gene expression: correlation between enhancer activity and gene expression across cell types, gene expression level of target genes, genomic distance between an enhancer and its target gene, enhancer signal, average gene activity in the region between the enhancer and target gene and enhancer–enhancer correlation. We downloaded enhancer annotations for 104 cell types from http://www.enhanceratlas.org/.

#### *Enhancer-gene correlation* (DNase-seq and microarray gene expression)

Sheffield *et al.* (2013) correlated DNase I signal and gene expression levels using data from 112 human samples representing 72 cell types to identify regulatory elements and to predict their targets^41^. We downloaded these predictions from http://dnase.genome.duke.edu/ and used the correlation as the quantitative score. Cell-type specific links were not reported.

#### JEME

Cao et al. (2017) computed correlations between gene expression and various enhancer features (*e.g.*, DNase1, H3K4me1) across multiple cell types to identify a set of putative enhancers^42^. Then a sample-specific model is used to predict the enhancer gene connections in a given cell type. We downloaded the lasso-based JEME predictions in all ENCODE+Roadmap cell types from http://yiplab.cse.cuhk.edu.hk/jeme/. We used the JEME confidence score as a quantitative score.

#### TargetFinder

Whalen et al. 2016 built a model to predict whether nearby enhancer-promoter pairs are located at anchors of Hi-C loops based on chromatin features^44^. We downloaded the TargetFinder predictions from https://raw.githubusercontent.com/shwhalen/targetfinder/master/paper/targetfinder/combined/output-epw/predictions-gbm.csv. For each distal element-gene pair in our dataset, we searched to see if the element and gene TSS overlapped an enhancer and promoter loop listed in this file. If so, we assigned the pair a score corresponding to the ‘prediction’ column from this file; otherwise the pair received a score of 0.

### Cell-type specific gene set enrichments

We assessed whether the cell-type specificity of the ABC predictions for IBD variants could aid in identifying gene pathways enriched in IBD GWAS loci. To do so, we defined 7 cell type categories based on the biosamples available in our compendium and based on biological categories relevant IBD: mononuclear phagocytes, B cells, T cells, other hematopoietic cells, fibroblasts, epithelial cells or tissues, and other cells or tissues. We then examined the extent to which the genes predicted by ABC in any cell type category, or in each individual cell type category, were enriched for gene sets from the Molecular Signatures Database^46^, as described above.

### Assessing pleiotropy across 72 traits

We defined two classes of genes: (i) Genes linked by ABC-Max to more than one trait via a single variant. To identify such genes, we counted cases where a given variant was linked by ABC-Max to the same gene for more than one trait (that is, the linked variant has PIP >= 10% for each trait, overlaps an enhancer in a biosample that is globally enriched for that trait, and links to the same gene with the maximum ABC score among genes in the locus). (ii) Genes linked to multiple traits through different variants. To identify such genes, we identified genes that were predicted by ABC-Max to be linked to at least two different traits by two different variants, where that gene was not linked to the same two traits by any single variant. (*i.e.*, a gene linked to two traits by each of two variants would not fit this criteria). Because some of the 72 traits show high genetic correlation, we repeated these analyses in a subset of 36 traits that were selected to show pairwise genetic correlation below a threshold (|*rg*| < 0.2), plus IBD. We observed similar effects in this subset of the data, where genes linked to multiple traits via different variants were more likely to have complex enhancer landscapes and large amounts of nearby noncoding genomic sequence.

### Assessing the effect of PPIF and e-PPIF on mitochondrial membrane potential

We synthesized a pool of 105 gRNAs including 40 negative control gRNAs, 9 gRNAs targeting the promoter of *PPIF*, and 5 gRNAs targeting the *PPIF* enhancer (Agilent Technologies, Inc.; see **Table S12**), cloned these gRNAs into Crop-seq-opti (Addgene #106280), and transduced THP1 cells at a multiplicity of infection of 0.3 to ensure most cells contained 1 gRNA integration.

For untreated and LPS-stimulated conditions, we plated 10M cells per replicate with 1 μg/mL doxycycline. After 44 hrs, we added 1 μg/mL LPS and harvested cells for staining 4 hrs later. For the PMA LPS condition, we plated 10M cells per replicate and added 1μg/mL doxycycline for 48 hrs. To differentiate into macrophage-like cells, we added fresh media with 20 ng/mL PMA and 1μg/mL doxycycline for an additional 24 hrs, confirming that cells adhered to the tissue culture plate. We washed out the PMA and added fresh media with 1μg/mL doxycycline and incubated cells for 45 hrs to recover and further differentiate cells. We then added 100 ng/mL LPS for 3 hrs, harvested cells, washed 3x with cold PBS, and proceeded to mitochondrial staining.

We stained cells with MitoTracker Red (200nM, Thermo Fisher, M7512) and MitoTracker Green (200nM, Thermo Fisher, M7514) according to the manufacturer’s protocol and sorted cells into 3 bins according to their ratio of MitoTracker Red (which stains mitochondria dependent on Δψ_m_) to MitoTracker Green (which stains mitochondria independent of Δψ_m_), excluding a small population of depolarized cells with very low Δψ_m_ (**Figure S8e**). We extracted genomic DNA and amplified and sequenced gRNAs from cells in each bin as previously described^24^.

We aligned and counted gRNAs in each bin as described above for FlowFISH experiments. For each gRNA, we summed counts across the two biological replicates. We then calculated the frequency fold-change in **Fig. 5d** and **Fig. S8b** by dividing gRNA reads per million by the mean value for negative-control gRNAs, and dividing values in each bin by the value for Bin 3.

## Data and Code Availability

Immune cell line ATAC-seq and H3K27ac ChIP-seq: NCBI GEO GSE155555 ABC predictions in 131 biosamples: www.engreitzlab.org/resources/ ABC code: https://github.com/broadinstitute/ABC-Enhancer-Gene-Prediction

## Acknowledgements

This work was supported by the Broad Institute (E.S.L.) and an NIH Pathway to Independence Award (K99HG009917 and R00HG009917 to J.M.E.). J.M.E. was supported by the Harvard Society of Fellows and the BASE Research Initiative at the Lucile Packard Children’s Hospital at Stanford University. The authors thank Larry Schweitzer, Matteo Gentili, Moshe Biton, Chris Smillie, Aviv Regev, Masahiro Kanai, Daniel Graham, Noam Shoresh, Steven Gazal, Brian Cleary, Ran Cui, Patricia Rogers, Vidya Subramanian, Gavin Schnitzler, Raj Gupta, Melina Claussnitzer, Nasa Sinnott-Armstrong, Tim Majarian, Alisa Manning, and members of the Lander Lab, Hacohen Lab, and Variant-to-Function Initiative for discussions or technical assistance. This research has been conducted using the UK Biobank Resource.

## Supplemental Figures

**Fig S1.**
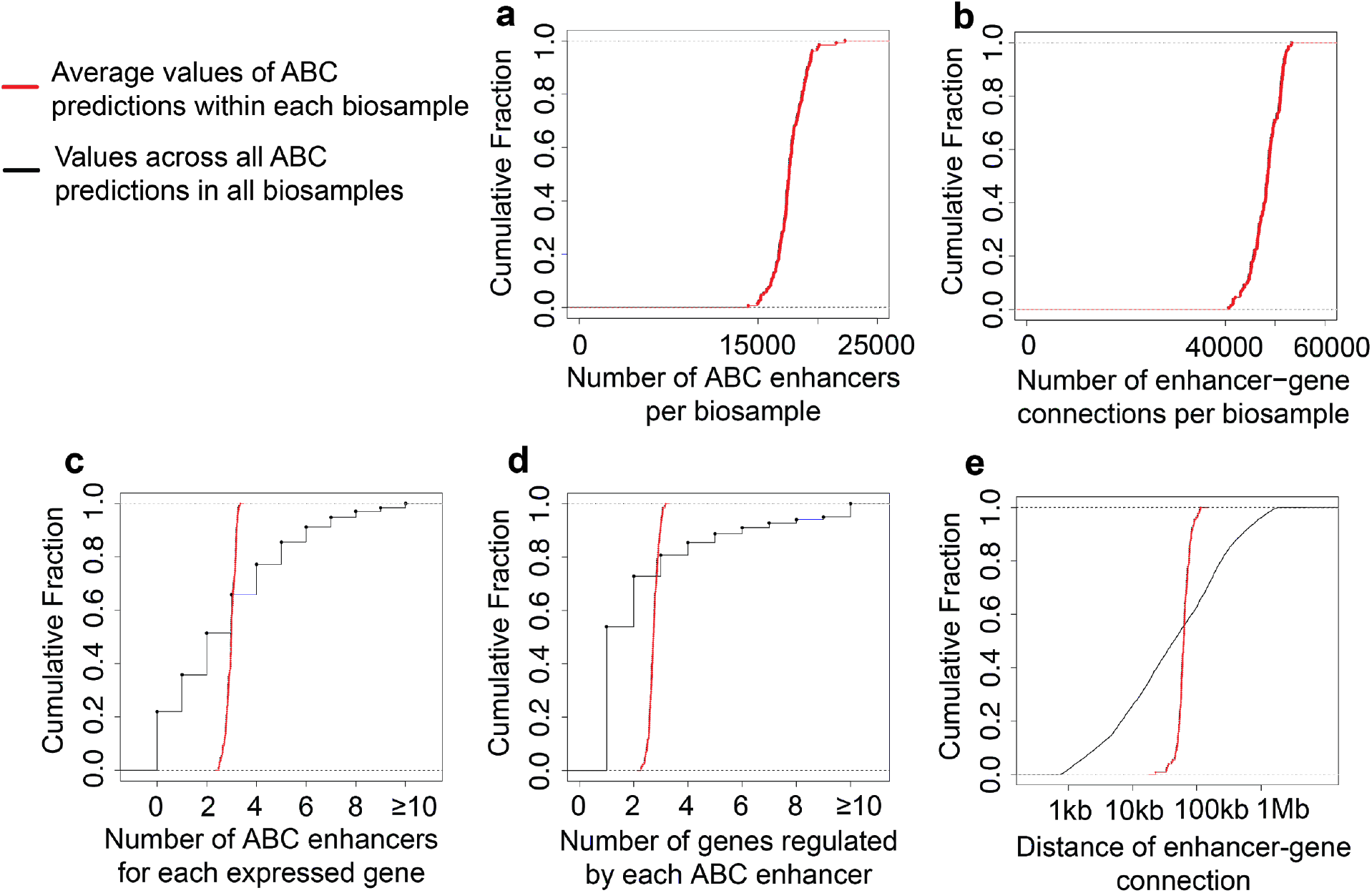
Properties of ABC Predictions. **(a)** Cumulative fraction of the number of ABC enhancers within each biosample (median = 17,605). **(b)** Cumulative fraction of the number of enhancer-gene connections within each biosample (median = 48,441). **(c)** Cumulative fractions of the number of enhancers predicted to regulate each gene across all biosamples (black line, median = 2, mean = 2.8) and the mean number of enhancers predicted to regulate each gene within each biosample (red line, median = 2.8). **(d)** Cumulative fractions of the number of genes regulated by each ABC enhancer across all genes and all biosamples (black line, median = 1, mean = 2.7) and the mean number of genes regulated by each ABC enhancer within each biosample (red line, median = 2.7). **(e)** Cumulative fractions of the genomic distances between the enhancer and the gene for each predicted enhancer-gene connection across all genes and all biosamples (black line, median = 62,929bp) and the median genomic distance between each enhancer-gene connection within each biosample (red line, median = 62,782 bp).

**Fig S2.**
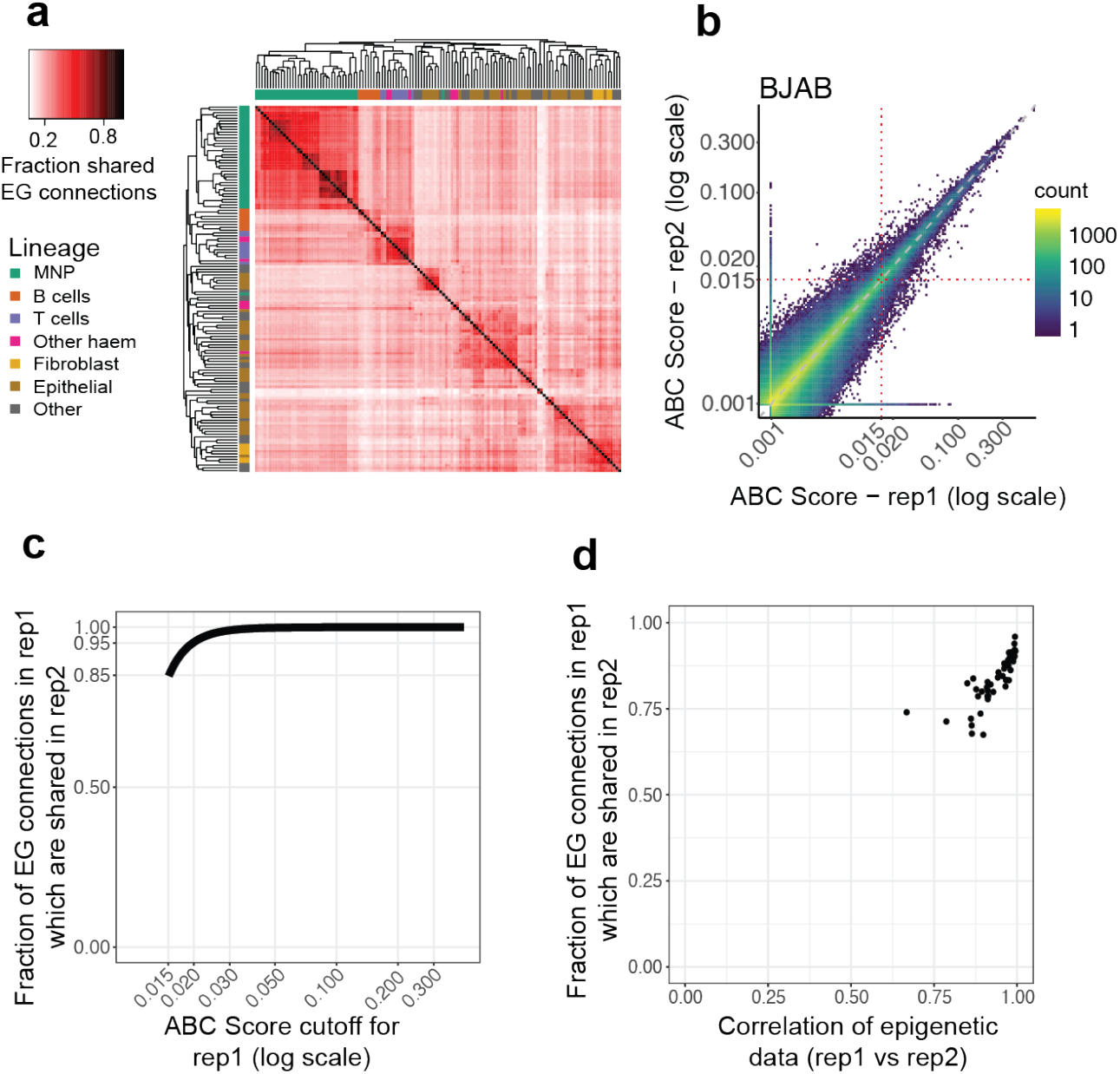
Distinctness and Reproducibility of ABC predictions. **(a) Distinctness of predictions across biosamples.** Biosample vs. Biosample (131 x 131) heatmap. The color of the (*i*,*j*) pixel in the heatmap represents the fraction of enhancer-gene (EG) connections in biosample *i* that have a corresponding overlapping prediction in biosample *j*. Two connections are considered overlapping if the predicted genes are the same and the enhancer elements overlap. Rows and columns are ordered by hierarchical clustering. A median of 19% (median of row medians) of enhancer-gene connections are shared across distinct biosamples. **(b) Quantitative reproducibility of ABC Predictions.** ABC Scores computed using independent biological replicates of epigenomic data (ATAC-Seq and H3K27ac ChIP-Seq) from the BJAB cell line. Each data point is an element-gene pair. **(c) Fraction of shared enhancer-gene connections between replicates increases as ABC Score cutoff increases.** X-axis: Cutoff on the ABC Score. Y-axis: For a given cutoff of the ABC Score, the fraction of element-gene pairs with an ABC score greater than the cutoff in replicate 1 that have an ABC score > 0.015 in replicate 2. Separate curves are computed for each biosample and then the average across biosamples is plotted. **(d) Fraction of shared enhancer-gene connections increases as reproducibility of underlying epigenetic data increases.** Each data point represents a biosample. X-axis: average correlation of ATAC-Seq (or DNase-Seq) and H3K27ac ChIP-Seq signal in candidate regions computed using replicate epigenetic experiments. Y-axis: Fraction of E-G connections with ABC Score > 0.015 in replicate 1 which also have ABC Score > 0.015 in replicate 2.

**Fig. S3.**
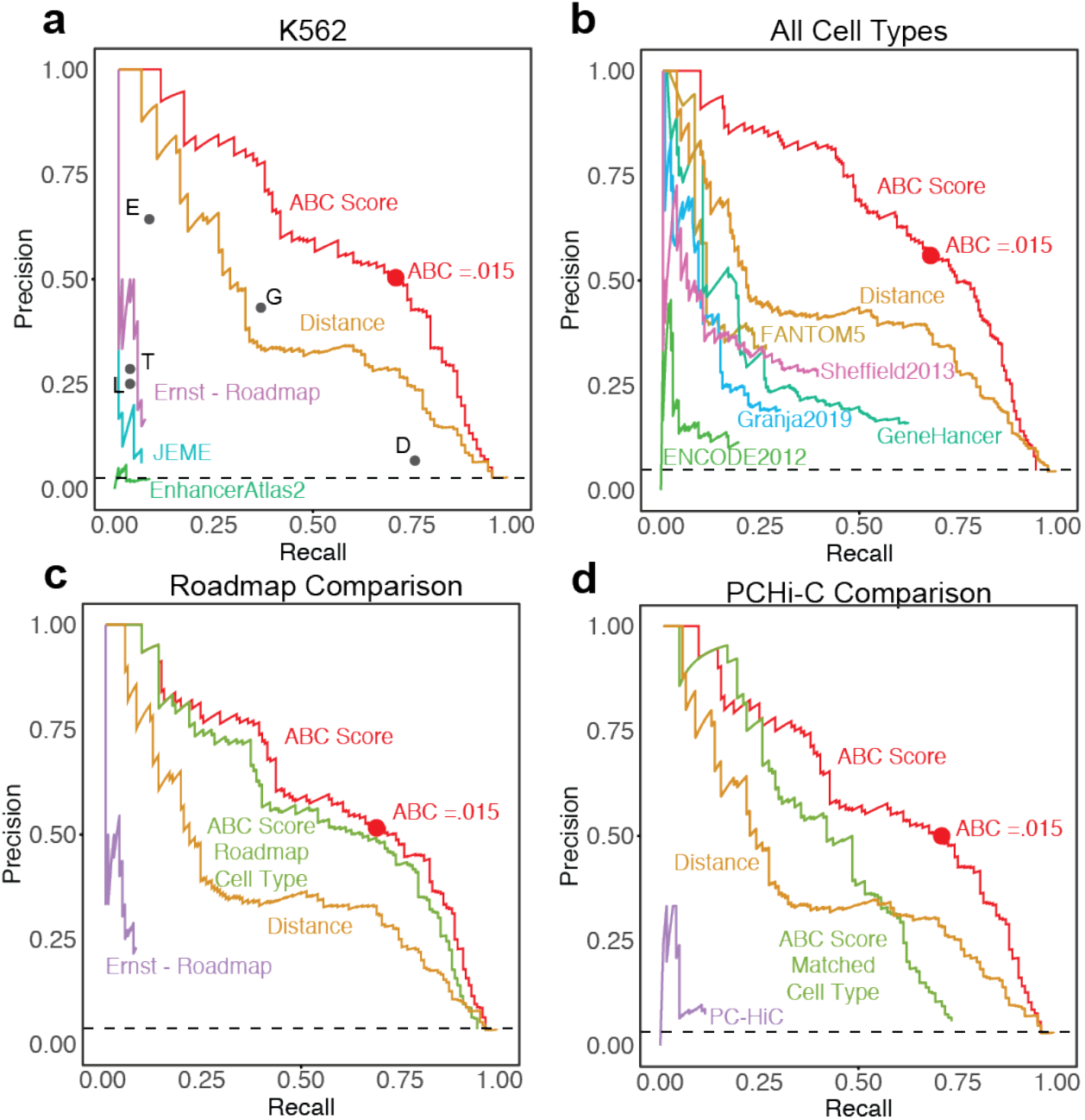
ABC performs well at identifying regulatory enhancer-gene connections in CRISPR datasets. **(a) Comparison of enhancer-gene predictors to experimental CRISPR data in K562 cells.** Each of these predictors makes K562-specific predictions. Curves represent continuous predictors. Dots represent binary predictors as follows: (E) Each gene is predicted to be regulated only by the element closest to its transcription start site, (G) each element is predicted to regulate only the nearest (to TSS) expressed gene, (T) TargetFinder method^44^, (L) elements and genes at opposite ends of HiCCUPS loops are predicted as a connection^89^, (D) each element is predicted to regulate each gene in the same contact domains^89^. Red dot on ABC score curve represents the precision and recall achieved using a threshold on the ABC score of 0.015. **(b) Comparison of ABC and other enhancer-gene predictors in full CRISPR dataset.** Comparison of enhancer-gene predictors to experimental CRISPR data in K562, GM12878, NCCIT, BJAB (+/− stimulation), Jurkat (+/− stimulation), THP1 (+/− stimulation) cells and primary hepatocytes. For ABC, we used the predictions in the cell type corresponding to the CRISPR experiments. Because ABC is the only method that makes predictions in all of these cell types, we used this plot to compare ABC to other methods that make predictions without cell-type information. We consider each enhancer-gene pair predicted by these methods to be a prediction in all cell types. Red dot on ABC score curve as in (a). **(c) Comparison of ABC and Ernst-Roadmap predictions.** Comparison of enhancer-gene predictors to experimental CRISPR data in K562, GM12878, and unstimulated Jurkat, BJAB, THP1 cells. Red line represents comparison of ABC scores derived using epigenetic data in the same cell type as the CRISPR experiment was performed. Red dot on ABC score curve as in (a). To compare Roadmap predictions to CRISPR data, we made cell type substitutions because the Roadmap predictions did not include BJAB, Jurkat, and THP1 cells: for BJAB CRISPR data we compared to predictions in the Roadmap B cell sample (E032); for THP1 data we used the Roadmap monocyte sample (E124); and for Jurkat data we used the Roadmap T cell sample (E034). To directly compare the performance of ABC and Ernst-Roadmap methods in matched cell types, we also calculated ABC performance using the same cell type substitutions (green line). **(d) Comparison of ABC to Promoter-Capture Hi-C.** Comparison of enhancer-gene predictors to experimental CRISPR data in K562 and unstimulated BJAB, THP1 and Jurkat cells. Red line represents comparison of ABC Scores derived using epigenetic data in the same cell type as the CRISPR experiment was performed. Red dot on ABC Score curve as in (a). To compare promoter-capture Hi-C predictions to CRISPR data, we made cell type substitutions because the these predictions did not include K562, BJAB, Jurkat, and THP1 cells: for K562 CRISPR data we compared to predictions in erythroblasts; for BJAB CRISPR data we compared to total B cells; for THP1 data we compared to monocytes; and for Jurkat data we compared to total CD4+ T cells. To directly compare the performance of ABC and PC-HiC methods in matched cell types, we also calculated ABC performance using the same cell type substitutions (green line).

**Figure S4.**
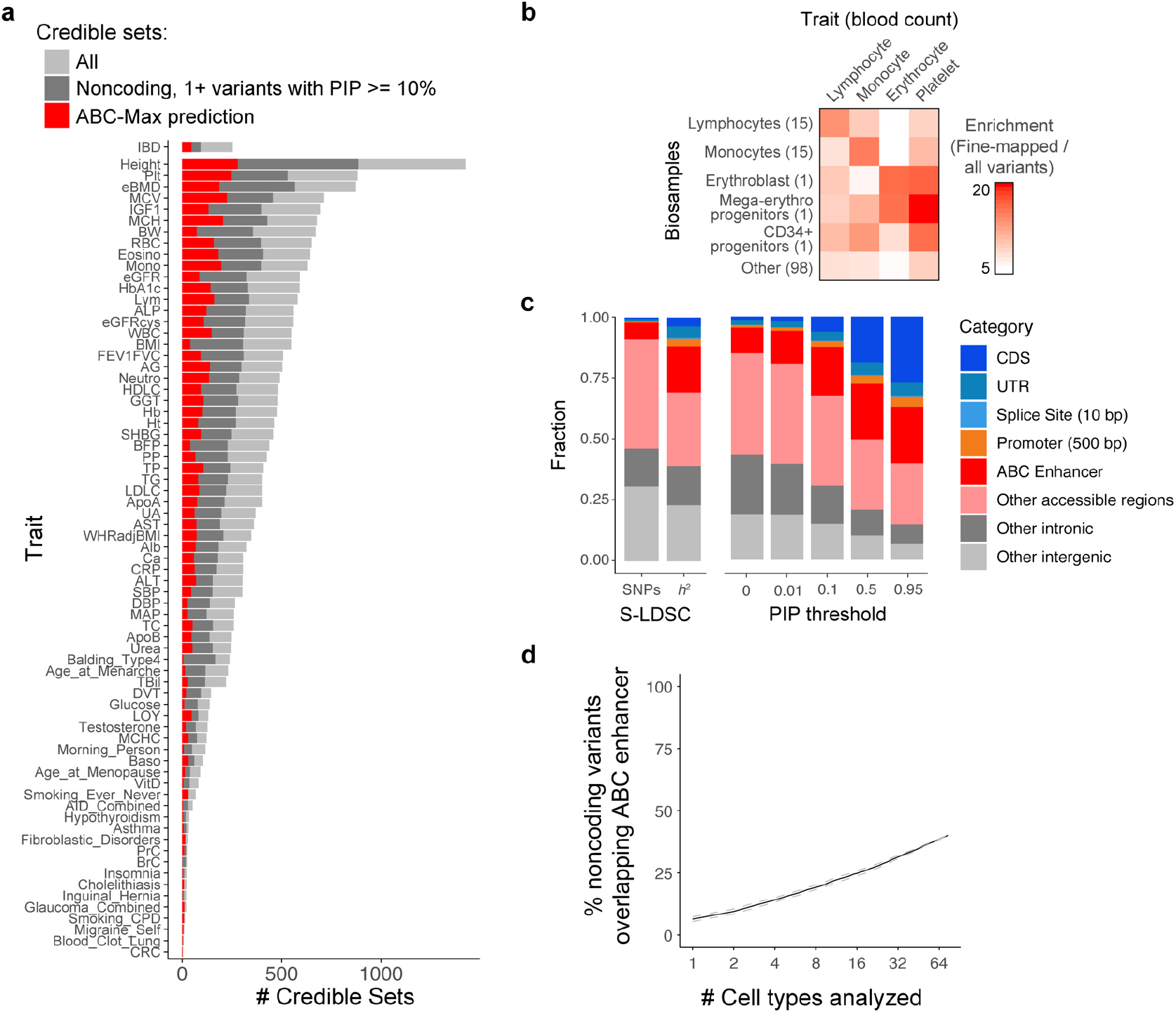
Fine-mapped GWAS variants are highly enriched in ABC enhancers. **(a)** Number of credible sets analyzed for 72 diseases and complex traits. Light gray shows total number of fine-mapped credible sets. Dark gray shows number of such credible sets with no coding or splice site variants, and at least one variant with PIP >= 10%. Red shows number of credible sets for which ABC-Max makes a prediction (*i.e.*, a variant with PIP >= 10% overlaps an ABC enhancer in a biosample that shows global enrichment for that trait). See **Table S7** for trait descriptions and additional statistics. **(b)** Enrichment of fine-mapped variants (PIP >= 10%) associated with 4 blood cell traits in ABC enhancers in the corresponding blood cell types or progenitors. Enrichment = (fraction of fine-mapped variants / fraction of all common variants) overlapping regions in each cell type. Numbers of biosamples in each category are shown in parentheses. **(c)** Fraction of variants or heritability for all 72 traits contained in different categories of genomic regions: coding sequences (CDS), untranslated regions (UTR), splice sites (within 10 bp of a intron-exon junction of a protein-coding gene), promoters (±250 bp from the gene TSS), ABC enhancers in 131 biosamples, other accessible regions not called as ABC enhancers, and other intronic or intergenic regions. In cases where a variant overlaps more than one category, the variant was assigned to the first category that it overlapped (*i.e.*, variants in coding sequences were not also counted as overlapping ABC enhancers, see Methods). Left: All common variants (1000 Genomes) or heritability (*h*^2^, as estimated by S-LDSC in inverse-variance weighted meta-analysis across 74 traits). Right: Fraction of variants above a threshold on the fine-mapping PIP. **(d)** % of noncoding variants across all traits that overlap an ABC enhancer in an enriched biosample, as a function of the number of cell types analyzed. Biosamples (131) were grouped into 74 cell types/tissues; and analyzed in random order. Black line: mean across 20 random orderings. Dashed gray lines: 95% confidence intervals.

**Figure S5.**
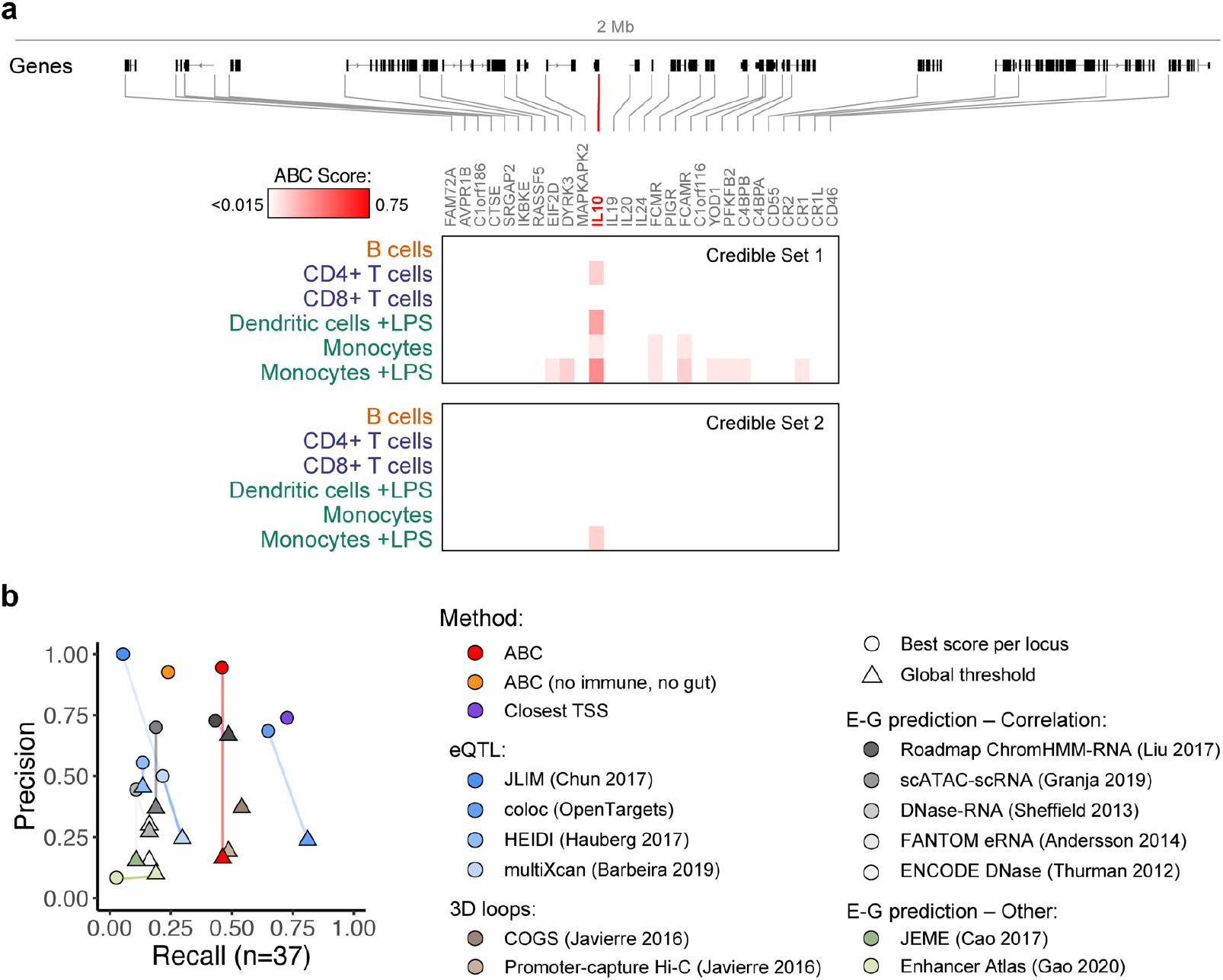
ABC enhancer maps connect GWAS variants to IBD genes. **(a)** ABC predictions for IBD credible sets linked to *IL10*. Heatmap shows ABC scores for each gene within 1 Mb in selected primary immune cell types. Credible Set 1 is linked by ABC to multiple genes, but *IL10* (red) has the strongest ABC score in any cell type. **(b)** Precision-recall plot for identifying known IBD genes, comparing additional variations on the prediction methods (superset of data in **Fig. 2a**). For ABC, we compared ABC-Max (assigning each credible set to the gene with the maximum ABC score, red circle), ABC-Max excluding all immune and gut tissue biosamples (orange circle), and ABC-All (assigning each credible set to all genes linked to enhancers, red triangle). For other methods that provided quantitative scores, we similarly compared choosing the gene with the best score per locus (circles) with choosing all genes above the global thresholds previously reported in each study (triangles). In most cases, the best gene per locus outperformed using a global threshold.

**Figure S6.**
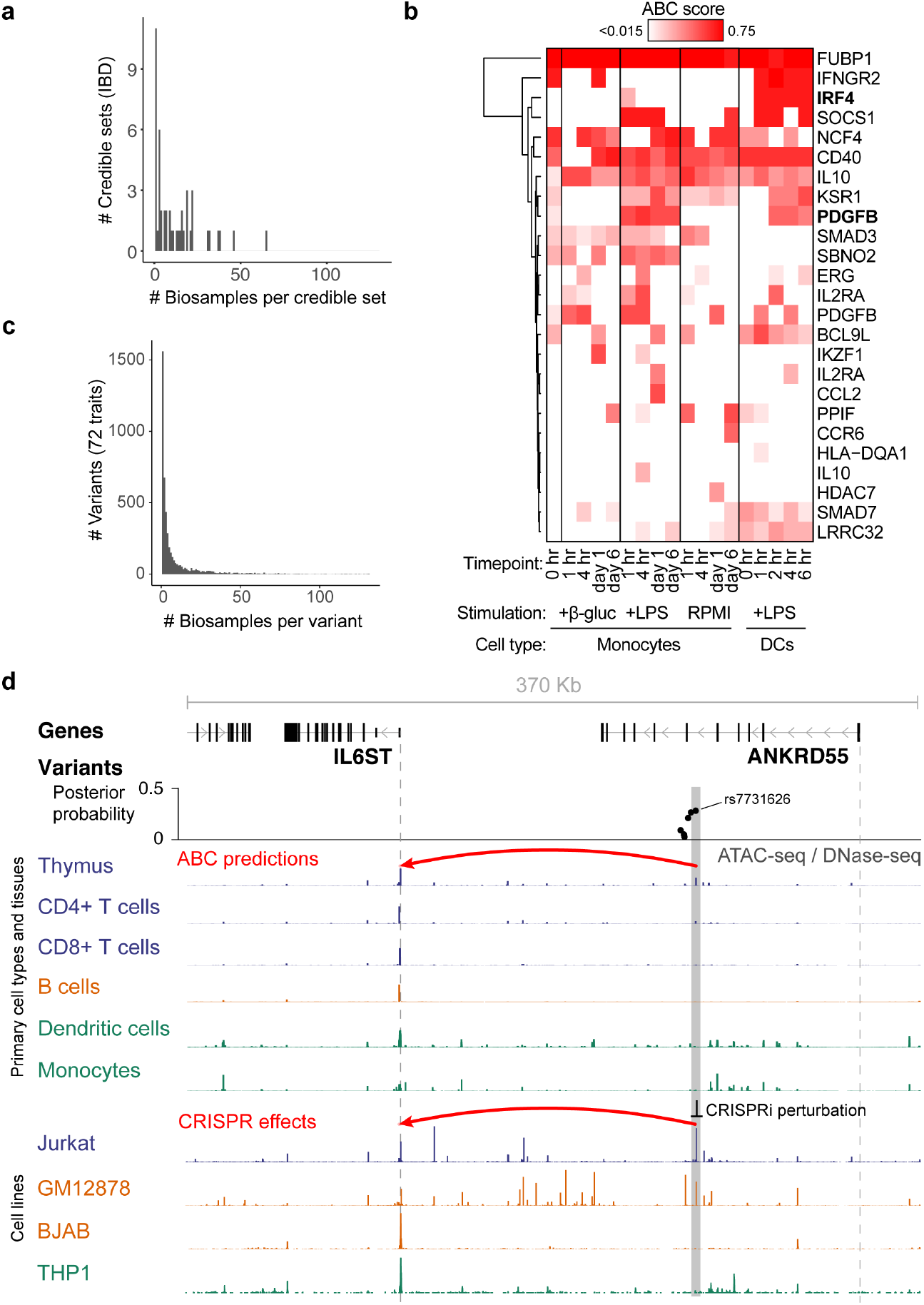
Cell-type specificity of ABC predictions. **(a)** Histogram of number of biosamples predicted by ABC-Max per noncoding IBD credible set (total = 47 credible sets). **(b)** Heatmap of ABC scores for predicted IBD genes in resting and stimulated mononuclear phagocytes (from epigenomic data in monocytes^90^ and dendritic cells^91^). IRF4 and PDGFB (bold) are two examples where ABC predictions are specific to a particular stimulated state (+LPS) and are not observed in unstimulated states. **(c)** Histogram of number of biosamples predicted by ABC-Max per variant (total = 4,976 unique variants across 72 traits). **(d)** A variant in an intron of *ANKRD55* is predicted by the ABC Model to regulate *IL6ST* in thymus. Gray bar highlights the variant overlapping the predicted ABC enhancer. Vertical dotted lines represent TSSs. Red arc at top denotes ABC-Max prediction. Rec arc at bottom denotes that CRISPRi of the highlighted enhancer significantly affects the expression of *IL6ST* only in Jurkat cells.

**Figure S7.**
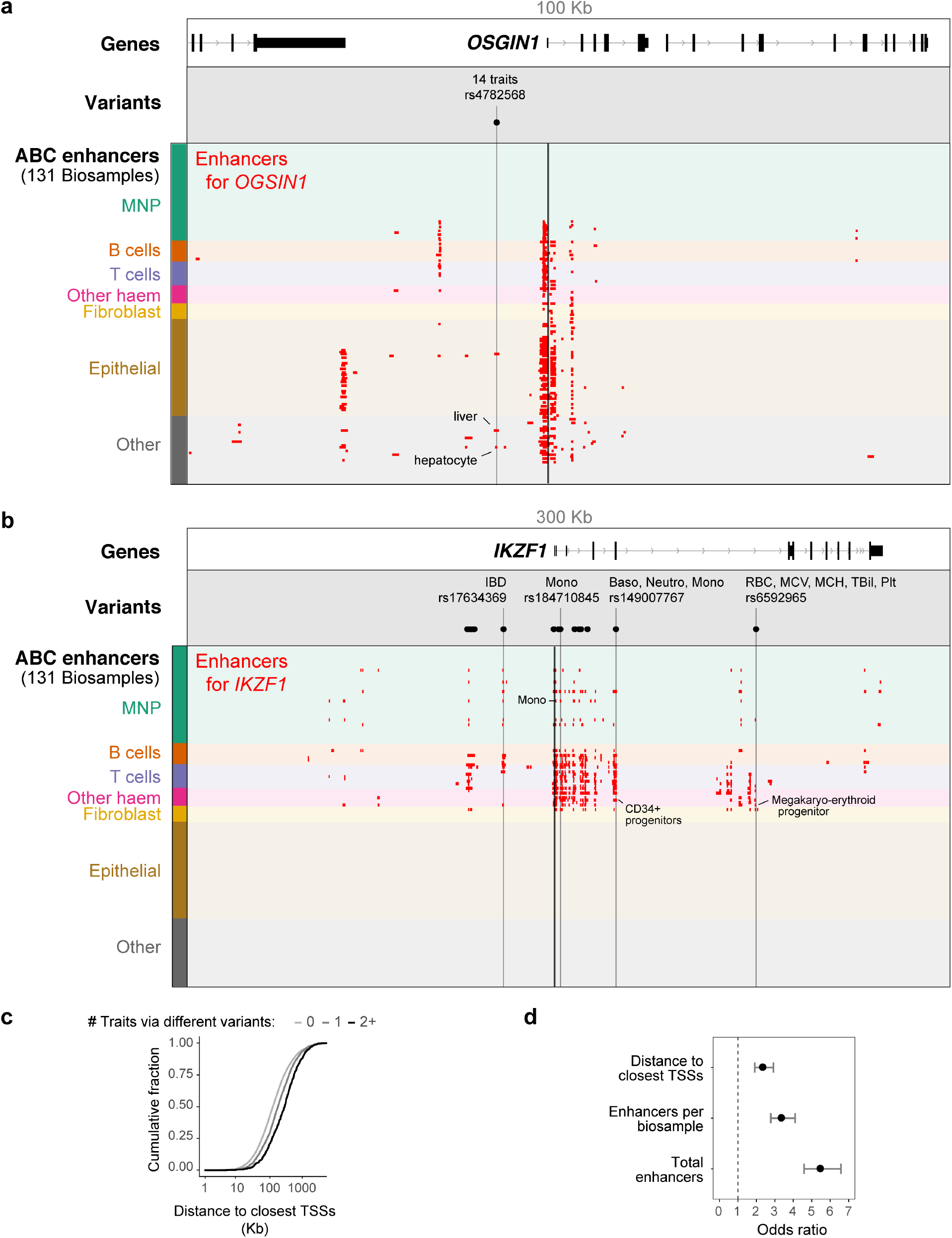
Genes linked by ABC to multiple traits. **(a)** ABC links *OSGIN1* to 14 traits via rs4782568. Red boxes mark enhancers predicted to regulate *OSGIN1*. Thick black line marks the *OSGIN1* TSS. **(b)** Same for *IKZF1* locus. Black dots mark fine-mapped noncoding variants (PIP >= 10%) associated with one or more traits linked to *IKZF1* by ABC-Max. **(c)** Distance between the TSSs of neighboring genes for each gene linked by ABC-Max to zero traits, one trait, or two or more traits through different variants. **(d)** The complexity of a gene’s enhancer landscape is correlated with the odds of the gene being linked to multiple GWAS traits. Wald odds ratios and 95% CIs for the genes in the top decile, compared to other genes, are shown for 3 gene-based metrics: the total number of enhancers linked to the gene by ABC in any biosample, the number of enhancers linked to a gene per biosample in which the gene’s promoter is active, and genomic distance to the nearest neighboring TSS on either side of the gene.

**Figure S8.**
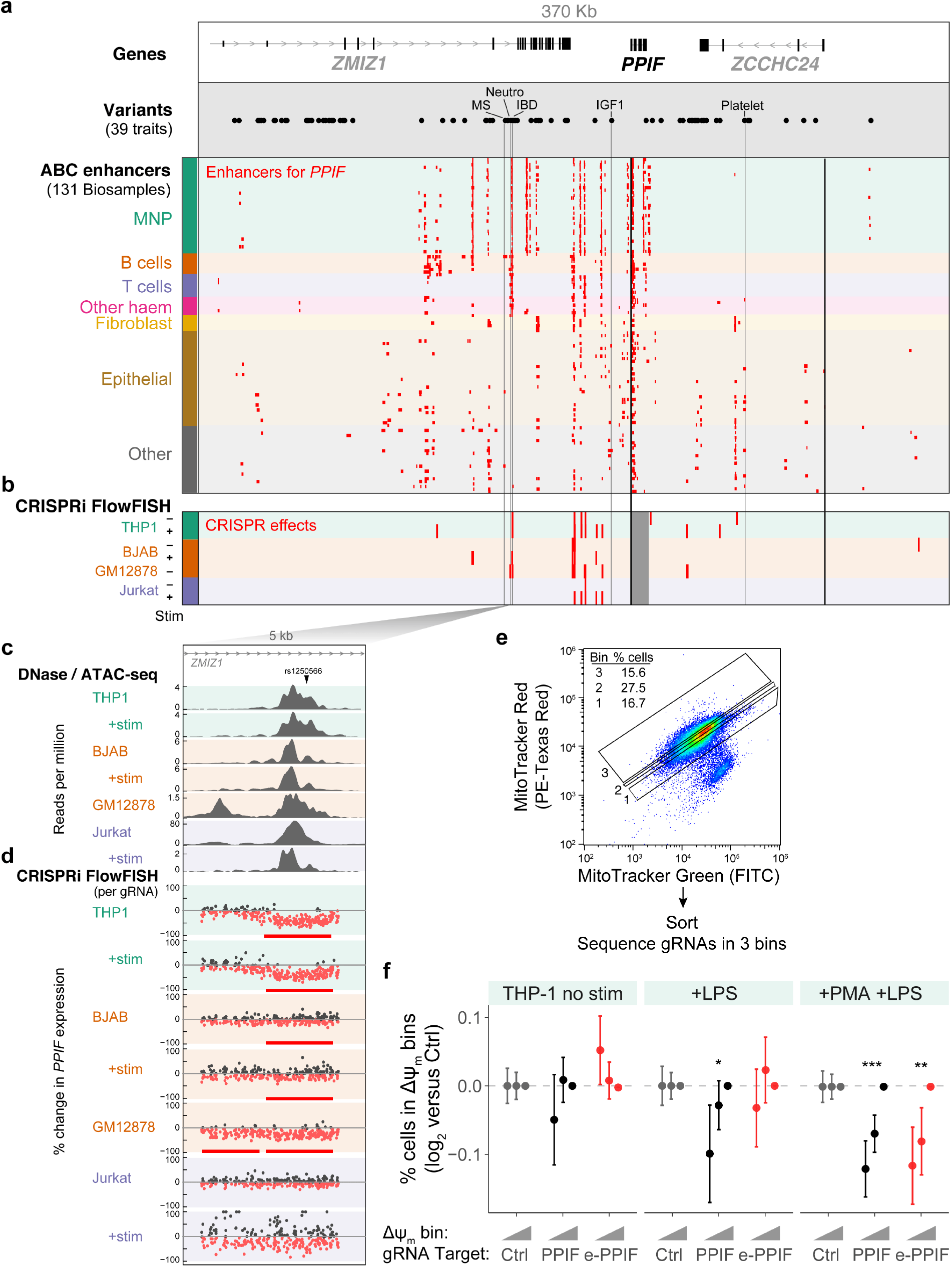
Enhancers and variants connected to *PPIF*. **(a)** ABC predictions for variants near PPIF. Black dots represent either (i) fine-mapped variants (PIP >= 10%) for IBD and UK Biobank traits, or (ii) lead variants for any phenotype from the GWAS Catalog^92^. “IBD” label points to rs1250566. Red boxes mark enhancers predicted to regulate PPIF. Thick back lines mark TSSs. Thin black lines mark selected variants. **(b)** CRISPRi-FlowFISH data for PPIF in 7 immune cell lines and stimulated states. Red boxes mark distal enhancers (CRISPR gRNAs lead to a significant decrease in the expression of PPIF). Dark gray box marks the gene body of *PPIF*, where CRISPRi cannot accurately assess the effects of putative regulatory elements^24^. **(c)** Chromatin accessibility in 5-kb regions around the PPIF enhancer and promoter. Signal tracks show ATAC-seq (for THP1 and BJAB) or DNase-seq (for GM12878 and Jurkat) data in reads per million. **(d)** Effect of each tested gRNA on PPIF expression, as measured by CRISPRi-FlowFISH (see Methods). Dots: gRNAs whose effect estimate is >0% (black) or <0% (red). Red bars show regions where gRNAs have a significant effect on gene expression (FDR < 0.05), as compared by a two-sided t-test to negative control gRNAs. **(e)** Schema of pooled CRISPRi screen to examine the effects of *PPIF* and *e-PPIF* on mitochondrial membrane potential (Δψ_m_). Cells expressing a pool of gRNAs were stained with MitoTracker Red and MitoTracker Green and sorted into 3 bins of increasing Red:Green ratios. gRNAs from cells in each bin were PCR-amplified, sequenced, and counted. **(f)** Effects of CRISPRi gRNAs (targeting e-PPIF, PPIF promoter, or negative controls (Ctrl)) on Δψ_m_, quantified as the frequency of THP1 cells carrying those gRNAs with low or medium versus high MitoTracker Red signal (corresponding to Bins 1, 2, and 3, respectively; superset of data in **Fig. 5d**). We tested THP1 cells in unstimulated conditions, stimulated with LPS, and differentiated with PMA and stimulated with LPS (see Methods). Error bars: 95% confidence intervals for the mean of 40, 9, and 5 gRNAs for Ctrl, PPIF, and e-PPIF, respectively. Two-sided rank-sum *P* < 0.05 (*), <0.01 (**), or <0.005 (**) versus Ctrl.

## Supplementary Tables

**Table S1 |** Epigenomic data collected in immune cell lines.

**Table S2 |** Metrics for ABC predictions in 131 biosamples.

**Table S3 |** CRISPRi-FlowFISH data: data per guide.

**Table S4** | CRISPRi-FlowFISH data: summary per candidate element.

**Table S5** | Comparison of CRISPR data to enhancer-gene predictions.

**Table S6** | Enrichment of GWAS variants in ABC enhancers across biosamples.

**Table S7** | Summary of diseases and traits.

**Table S8** | ABC predictions for IBD GWAS loci.

**Table S9** | ABC-Max predictions for 72 diseases and complex traits.

**Table S10** | References linking predicted genes to effects on experimental colitis.

**Table S11** | ABC and ABC-Max metrics for all genes.

**Table S12** |*PPIF* and mitochondrial membrane potential: CRISPRi data per guide.

